# Differential immune- and apoptosis-related gene signatures in pancreatic alpha and beta cells contribute to their fate in type 1 diabetes

**DOI:** 10.1101/2025.07.26.666935

**Authors:** Xiaoyan Yi, Eugenia Martin-Vazquez, Sayro Jawurek, Priscila L. Zimath, Junior Garcia Oliveira, Jose Maria Costa-Junior, Erwin Ilegems, Johnna D. Wesley, Alexandra C. Title, Burcak Yesildag, Decio L. Eizirik

**Affiliations:** ULB Center for Diabetes Research, Medical Faculty, Université Libre De Bruxelles (ULB), Brussels, Belgium; InSphero AG, Schlieren 8952, Switzerland; Novo Nordisk Research Center, Novo Nordisk A/S, 2760 Maaloev, Denmark; Novo Nordisk US R&D, Lexington, MA, 02128 United States

## Abstract

Both alpha and beta cells are dysfunctional in type 1 diabetes (T1D), but beta cells die while alpha cells survive the immune attack. Understanding the mechanisms underlying alpha-cell resistance could identify new approaches to protect beta cells. Herein, we analysed single-cell datasets from human alpha and beta cells under basal/unstimulated conditions and under immune-mediated stress. Alpha cells exhibit enhanced immune-like gene expression compared to beta cells. We also found that the tumour suppressor Maternally Expressed Gene 3 (*MEG3*), a T1D risk gene, is highly expressed in beta cells while almost undetectable in alpha cells. These observations were confirmed by analysing bulk RNA-sequencing data from fluorescence-activated cell-sorted alpha and beta cells isolated from primary human islets from non-diabetic donors. Additionally, *MEG3* knockdown in human insulin-producing EndoC-βH1 cells and human islets microtissues decreased cytokine-induced damage and apoptosis, preserving beta-cell function under inflammatory conditions. The fact that alpha cells exhibit increased immune-like and anti-apoptotic activity as compared to beta cells suggests that they are better equipped to endure the autoimmune assault in T1D. In addition, the marked difference in the expression of the pro-apoptotic factor *MEG3* in beta cells compared to alpha cells may explain, at least in part, why beta cells are more susceptible to damage and cell death in a diabetogenic environment than neighbour alpha cells within the same islet.

## INTRODUCTION

Type 1 diabetes (T1D) is characterized by both islet dysfunction and autoimmune-mediated loss of beta cells (1, 2). More than 90 genes have been identified as “T1D candidate genes” (3) and >80% of these genes are expressed in human beta cells. Furthermore, exposure to pro-inflammatory cytokines modifies human islet function and gene expression (4), suggesting that beta cells-in dialog with the immune system-play an important role in their own autoimmune-mediated destruction.

In T1D the immune system destroys the insulin-producing beta cells but not the neighbouring glucagon-producing alpha cells, despite both cell types being dysfunctional (5, 6). Analysis of single-cell RNA sequencing (scRNA-seq) of human induced pluripotent stem cell (hiPSC)-derived islet-like cells exposed to IFN-α (a major player in the early stages of T1D (7–9) demonstrated important differences between alpha and beta cells (10, 11). Thus, the genes that encode the heavy chain of the HLA class I molecule (*HLA-A, -B* and *-C*) are increased in IFN-α-treated alpha cells to a similar extent as in beta cells (10), but the atypical *HLA-E*, a protective molecule against the autoimmune assault (8), is upregulated in alpha compared to beta cells. These findings were subsequently confirmed in native human islets from individuals with T1D (12). Moreover, IFN-α-induces higher upregulation of Bcl-2-like protein 1 (*BCL2L1*), an inhibitor of intrinsic apoptosis pathway, and of *BiP*, an inhibitor of ER stress, in alpha cells than in beta cells (10).

Understanding how alpha cells can endure the protracted immune assault associated with T1D could provide useful information and new therapeutic targets to protect beta cells and limit or prevent the progression of T1D. In the current study, we analysed six single-cell and one bulk RNA-seq datasets of human islets. The data is comprised of non-diabetic donors; donors positive for one or two autoantibodies (AAB1+/2+); T1D-positive donors; individuals with type 2 diabetes (T2D) (13); and datasets from islet-like cells differentiated *in vitro* from stem-cells (10, 14, 15). Our bioinformatic analysis revealed that alpha cells consistently exhibited enhanced immune-like activity as compared to beta cells under basal conditions and in T1D. Additionally, we identified a potential new mediator of beta cell susceptibility to cytokines, namely Maternally Expressed Gene 3 (*MEG3*). The expression of this gene was markedly increased in beta compared to alpha cells among multiple datasets and biological samples. *MEG3* is a long non-coding RNA (lncRNA) highly expressed in brain and placenta (16, 17). Genome-wide association studies identified a single nucleotide polymorphism within the imprinted DLK1-MEG3 region on chromosome 14q32.2 associated with T1D risk (18) but the role of *MEG3* in human islet cells remains unknown. Using the human insulin-producing beta-cell line EndoC-βH1 (19) and native re-aggregated human islets microtissues (hIsMTs) (20), we presently evaluated the impact of *MEG3* on inflammatory stress and cell death triggered by pro-inflammatory cytokine exposure *in vitro* (1).

## MATERIALS AND METHODS

### Collection of raw scRNA-seq data

For the human islets datasets, raw scRNA-seq FASTQ files (10X Genomics) were downloaded from the Human Pancreas Analysis Program (HPAP) portal (https://hpap.pmacs.upenn.edu) (**Table S1**). HPAP-young and HPAP-older donors represent matched T1D and T2D controls curated by the HPAP consortium based on similar age and body mass index (BMI) characteristics. We analyzed 15 HPAP-young donors (HPAP-022, HPAP-026, HPAP-027, HPAP-034, HPAP-035, HPAP-036, HPAP-037, HPAP-039, HPAP-040, HPAP-044, HPAP-047, HPAP-056, HPAP-082, HPAP-099 and HPAP-104), 13 HPAP-older donors (HPAP-052, HPAP-053, HPAP-054, HPAP-059, HPAP-063, HPAP-074, HPAP-075, HPAP-077, HPAP-080, HPAP-093, HPAP-101, HPAP-103 and HPAP-105), 9 AAB1+ donors (HPAP-024, HPAP-029, HPAP-038, HPAP-045, HPAP-049, HPAP-050, HPAP-072, HPAP-092 and HPAP-114), 2 AAB2+ donors (HPAP-043 and HPAP-107), 11 T1D donors (HPAP-020, HPAP-021, HPAP-023, HPAP-032, HPAP-055, HPAP-064, HPAP-071, HPAP-084, HPAP-087, HPAP-113 and HPAP-123) and 10 T2D donors (HPAP-051, HPAP-057, HPAP-058, HPAP-061, HPAP-065, HPAP-070, HPAP-079, HPAP-083, HPAP-085 and HPAP-088). The clinical characteristics of these donors are available at the HPAP portal (https://hpap.pmacs.upenn.edu/explore/download?donor) by referencing their respective donor IDs. The raw scRNA-seq FASTQ files (10X Genomics) of hiPSC-derived and human embryonic stem cells (hESC)-derived islet-like cells were retrieved from the Gene Expression Omnibus (GEO) database under the accession numbers: GSE190726 (Chandra et al., hESC stage 6), GSE203384 (Szymczak et al., iPSC stage 7), GSE20083 (hESC-derived cells matured retrieved *in vivo* after transplantation into humanized mouse) and GSE20084 (hESC-derived cells differentiated *in vitro*), as summarized in **Table S1**.

### scRNA-seq data processing

Raw scRNA-seq data (10x Genomics) were processed using Cell Ranger (v6.1.2) (21) with default parameters. The cellranger count pipeline was used to perform read alignment to the human reference genome (GRCh38), barcode and UMI quantification, and generation of the gene-by-cell expression matrix. Ambient mRNA contamination was removed using SoupX v1.6.1 (22), leveraging marker genes (*INS, GCG, SST, TTR, IAPP, PYY, KRT19* and *TPH1*) representative of the major cell types identified in the initial clustering of each sample. The corrected gene count matrix was then imported into Seurat v4.3.0 (23) for downstream quality control. Cells were retained based on the following criteria: 1) genes expressed in at least three cells, and each cell expressing at least 200 genes; 2) potential doublets identified using scDblFinder v1.12.0 (24) were removed; 3) cells with fewer than 9 000 detected genes and fewer than 10 000 total UMI counts were kept; and 4) cells with >5% mitochondrial gene expression were excluded. Additional information is provided in Supplementary Methods.

### Bulk RNA-seq analysis processing

Raw FASTQ sequencing reads for the FACS-sorted alpha-cell and beta-cell samples from human islets were downloaded from the Human Pancreas Analysis Program (HPAP) data portal (https://hpap.pmacs.upenn.edu/) (**Table S1**). All samples underwent quality control using fastp v0.19.6 (25) and were subsequently analysed as described in Supplementary Methods.

### Gene Set Enrichment Analysis (GSEA)

Gene set enrichment analysis (GSEA) was performed using the fgsea v1.20.0 package (26) in R. The ranked gene list was generated based on the log2 fold change from differential expression analysis of both single-cell and bulk RNA-seq data. Gene sets of Reactome were downloaded from the Molecular Signatures Database (MSigDB v7.2, https://data.broadinstitute.org/gsea-msigdb/msigdb/release/7.2/) in GMT format. For each analysis, enrichment scores and associated statistics were calculated using the fgsea() function with the following parameters: minSize = 15, maxSize = 500, and nperm = 10,000. P-values were adjusted using the Benjamini-Hochberg method. Pathways with adjusted P < 0.05 were considered significantly enriched. A custom R script was used to visualize the top 15 significantly up- and downregulated pathways in each analysis, with pathway ranking based on normalized enrichment score (NES) and plotting implemented using ggplot2. If less than 15 pathways were significantly enriched (adjusted P-value < 0.05) in either direction, all significant pathways were displayed.

### Rank-rank Hypergeometric Overlap (RRHO) analysis

To compare the global transcriptomic similarity between HPAP-young (type 1 diabetes control) and HPAP-older (type 2 diabetes control) in the context of alpha-cell versus beta-cell differences, we performed Rank-Rank Hypergeometric Overlap (RRHO) analysis. We applied an enhanced RRHO pipeline, RedRibbon (27), which offers improved speed and accuracy compared to the original RRHO method (28), enabling efficient identification of shared up- or downregulated genes between two independent datasets (27). Detailed information on the procedure is provided in Supplementary Methods.

### EndoC-βH1 cell line culture and transduction

The human pancreatic beta cell line EndoC-βH1 was kindly provided by R. Scharfmann (Institut Cochin, Université Paris, Paris, France) (19). EndoC-βH1 cells were cultured in Dulbecco’s modified Eagle medium with 5.6 mM glucose (Gibco, Thermo Fisher Scientific) as previously described (29). All the results shown for EndoC-βH1 cells refer to independent biological samples, i.e., different passages. *MEG3* gene expression was silenced using 3 different siRNAs [Thermo Fisher Scientific]; siMEG3 #1 (siRNA n272552): 5′-GUGUUCACCUGCUAGCAAAtt-3′ and siMEG3 #2 (siRNA n272559): 5′-UCUUAUUUAUUCUCCAACAtt-3, both targeting 15 different transcript variants at exon 3 (si*MEG3* #1 at position 460; siMEG3 #2 at position 883), and siMEG3 #3 (siRNA n503814): 5′-GAACUGCGGAUGGAAGCUGtt-3′], targeting 15 transcript variants at exons 4 to 7 (positions ranging between 978 and 1342). AllStars Negative Control siRNA (siCTRL) (Qiagen) was used as a negative control; the siRNA control does not interfere with beta-cell gene expression, function or viability (30). Cells were transfected using Lipofectamine RNAiMAX lipid reagent (Invitrogen, Life Technologies) in Opti-MEM (Gibco, Thermo Fisher Scientific) with 30 nM of each siRNA, overnight as previously described (10). After transfection, cells were kept in culture for a 24h recovery period and subsequently exposed or not to human IFN-α (2000 U/ml; PeproTech), IFN-γ (1000 U/ml; PeproTech) alone or together with IL-1β (50 U/ml; PeproTech) for 24 or 48h. Cells were also exposed to the ER stressor Thapsigargin (1μM) for 48h. These conditions are based on our previously published time- and dose-response experiments (7, 10, 31).

### Human islets microtissues (hIsMTs) culture and transduction

Human islet microtissues (hIsMTs) (InSphero, MT-04-002-01-60) were generated after dispersion and reaggregation of primary human islets from a single donor (UNOS ID #ALIR128; male; 41y.o.; Hispanic; BMI 27.77; positive serologies: CMV IgG, EBV IgG; HbA1c: 5.3%; cause of death: head trauma). HIsMTs were transduced with AAVs encoding specific shRNA targeting *MEG3* (siMEG3 #1 sequence) from Vector Biolabs during aggregation for 5 days in Akura hanging drop plates (InSphero, CS-PF24) and then released into Akura 96-well plates (InSphero, CS-09-001-00). hIsMTs were cultured in standard culture medium (InSphero, CS-07-005-01) or standard culture medium containing cytokines: TNF-α (25 ng/mL, R&D Systems, 10291-TA-050); IL-1β (5 ng/mL, Sigma, H6291-10UG); IFN-γ (25 ng/mL, R&D Systems, 10067-IF-100) and IFN-α (10 ng/mL, Peprotech, 300-02AA) starting at day 9 after reaggregation, for 6 days. Medium was exchanged and cytokines were re-dosed every 2-3 days throughout the experiment.

### Cell viability assessments

EndoC-βH1 cell viability was determined by fluorescence microscopy using the nuclear dyes propidium iodide (10 μg/ml; Sigma-Aldrich) and Hoechst 33342 (10 μg/ml; Sigma-Aldrich), as previously described (30, 32). A minimum of 500 cells was counted per condition. Viability was evaluated by two independent observers, one of them being unaware of the sample identity. The agreement between the two observers was >90%. The results are expressed as percentage of apoptosis, calculated as the number of apoptotic cells/total number of cells and presented as fold change compared to siCTRL cytokines-treated cells. In EndoC-βH1 and hIsMTs, Caspase 3/7 activity was assessed using the Caspase-Glo 3/7 Assay System (Promega, G8091). Caspase reagent was added to wells containing cells or just culture medium (used as background control) and total Caspase 3/7 activity was evaluated by luminescence following manufacturer’s instructions.

### Quantitative real-time PCR, protein extraction and Western blot analysis

EndoC-βH1 were washed with PBS, detached and collected in lysis buffer. Poly(A) + RNA was isolated using the Dynabeads mRNA DIRECT Kit (Invitrogen) and reverse-transcribed using the Reverse Transcriptase Core Kit (Eurogentec), following the manufacturer’s protocol. HIsMTs were pooled (6 per well), washed twice in PBS and transferred to PCR-clean 2 mL Eppendorf tubes. QIAzol lysis buffer (Qiagen, 79306) was added and hIsMTs were vortexed and sonicated to aid thorough lysis. Additional details and primers composition are provided in Supplementary Methods.

Total protein was extracted using Laemmli buffer supplemented with phosphatase and protease inhibitors (Roche) and separated on 10% SDS– PAGE. The nitrocellulose membranes were probed using specific primary antibodies: Phospho-Stat1 (Tyr701) (58D6) Rabbit mAb #9167, Stat1 (9H2) Mouse mAb #9176 or Human/Mouse/Rat GAPDH Antibody R&D Systems # 2275-PC-100 diluted 1:1000 in TBST (TBS, 0.1% Tween 20) with 5% BSA (GAPDH was diluted 1:10 000). The densitometric values were quantified by ImageLab software version 6.1 (Bio-Rad Laboratories, RRID:SCR_014210) and normalized to GAPDH or the respective total protein form, after background subtraction. Full length uncropped original western blots used are provided in the Supplementary Material.

### Functional analysis and measurements of Insulin / CXCL10 secretion by ELISA

Glucose-stimulated insulin secretion was performed in hIsMTs sequentially in KRHB containing 2.8 mM and 16.7 mM glucose for 2h. ELISA (Human Insulin, Mercodia, Uppsala, Sweden for EndoC-βH1 cells, STELLUX® Chemi Human Insulin ELISA (Alpco, 80-INSHU-CH01) for hIsMTs or Human CXCL10/IP-10 Immunoassay, Quantikine ELISA kit, R&D Systems for both cell models) was used to quantify insulin or CXCL10 secretion to the culture medium (by 50 000 EndoC-βH1 cells / 6 hIsMTs / 200 μl of culture medium) as indicated in the figure legends. The cell culture supernatants were collected at the end of the experiments and immediately stored at −80°C until samples’ processing. For insulin secretion, all samples were diluted 10-fold. For CXCL10 secretion, samples under non-treated conditions were diluted 2-fold and samples exposed to cytokines were diluted 70-fold. The assay procedures and calculation of the results were conducted following the manufacturer’s recommendation.

### 3D staining and imaging

HIsMTs were washed twice in PBS (with Mg ^+^Ca ^+^), fixed for 15 min in 4% PFA, washed twice more with PBS and kept in PBS with 0.05% sodium azide until staining and imaging as described in Supplementary Methods.

### Quantification and Statistical Analysis

Data are presented as mean±SEM. One-way ANOVA was used as described in the figure legends. P values less than or equal to 0.05 were considered statistically significant. Statistical analyses were performed using GraphPad Prism software version 8 (GraphPad Software, La Jolla, USA). Statistics analysis for single-cell and bulk RNA-seq data are described in the corresponding sections above.

## RESULTS

### Single-cell RNA-seq (scRNA-seq) analysis of alpha and beta cells from primary and stem cell-derived human islets

We analyzed publicly available scRNA-seq data from 28 non-diabetic and autoantibody-negative human islet donors provided by the HPAP (**Table S1**). The non-diabetic data included 15 young, non-diabetic donors assigned as ‘T1D controls’ (HPAP-young) and 13 older, non-diabetic donors designated as ‘T2D controls’ (HPAP-older), based on their age and body max index (BMI). We also analyzed 11 autoantibody-positive donors: 9 positive donors for one islet autoantibody (HPAP-AAB1+) and 2 positive donors for 2 or more islet autoantibodies (HPAP-AAB2+). Additionally, samples from 10 individuals with T2D (HPAP-T2D) were also included in the analyses. We then re-analyzed two hiPSC-derived islet-like cell datasets: one from cells at stage 6 of differentiation, exhibiting an immature phenotype (identified as “Chandra et al., iPSC stage 6”, **Table S1**) and the other from cells at stage 7, which is the last stage of the hiPSC-to-islet-like cells differentiation protocol (10, 33, 34) (identified as “Szymczak et al., iPSC stage 7”, **Table S1**). Finally, we analyzed two human hESC-derived islet-like cell datasets: one from *in vivo* matured cells after transplantation into humanized mouse models (“Sintov et al., hESC *in vivo*”, **Table S1**) and the other from *in vitro* differentiated cells (“Sintov et al., hESC *in vitro*”, **Table S1**).

After stringent quality filtering, we obtained a total of 142 000 cells: 106 000 alpha cells and 36 000 beta cells for subsequent analysis (**Figure S1A**). For basal alpha cells, the largest contributor was the HPAP-older dataset, while for basal beta cells, the Szymczak et al. iPSC stage 7 dataset contributed with the highest number of cells (**Figure S1B**). Among alpha and beta cells from AAB1+, AAB2+, T1D, and T2D donors, T1D samples were associated with the largest proportion of alpha cells, while AAB1+ contributed with the largest proportion of beta cells (**Figure S1C-D**).

Under basal conditions, the alpha- and beta-cell populations derived from primary human islets exhibited well-defined cellular identities (**Figure 1A-B**). In the case of the hiPSC-derived islet-like cells, those at stage 7 displayed a more differentiated state compared to those at stage 6, as expected (**Figure 1C-D**). Alpha and beta cells derived from hESCs following transplantation and *in vivo* maturation showed a more mature cell identity as compared to their counterparts differentiated exclusively *in vitro* (**Figure 1E-F**). Additionally, we analyzed alpha- and beta-cell populations by stratifying donors from the HPAP into two groups: 1) a T1D-related group combining basal (young donors), autoantibody positive (AAB1+/2+) and T1D donors (**Figure S2A**) and 2) a T2D-related group combining basal (older donors) and T2D donors (**Figure S2B**). This analysis was performed to enable direct comparisons between alpha and beta cells within each disease setting in subsequent analyses. Next, to assess the global transcriptomic similarity between the HPAP-young and HPAP-older datasets in terms of basal alpha-cell versus beta-cell differences, we applied Rank-Rank Hypergeometric Overlap (RRHO) analysis using the RedRibbon tool (27). These analyses revealed a near-perfect correlation in both the up- and downregulated quadrants (**Figure S3A-B**). These results indicated that age has a modest impact on the transcriptional differences between alpha and beta cells.

**Figure 1.**
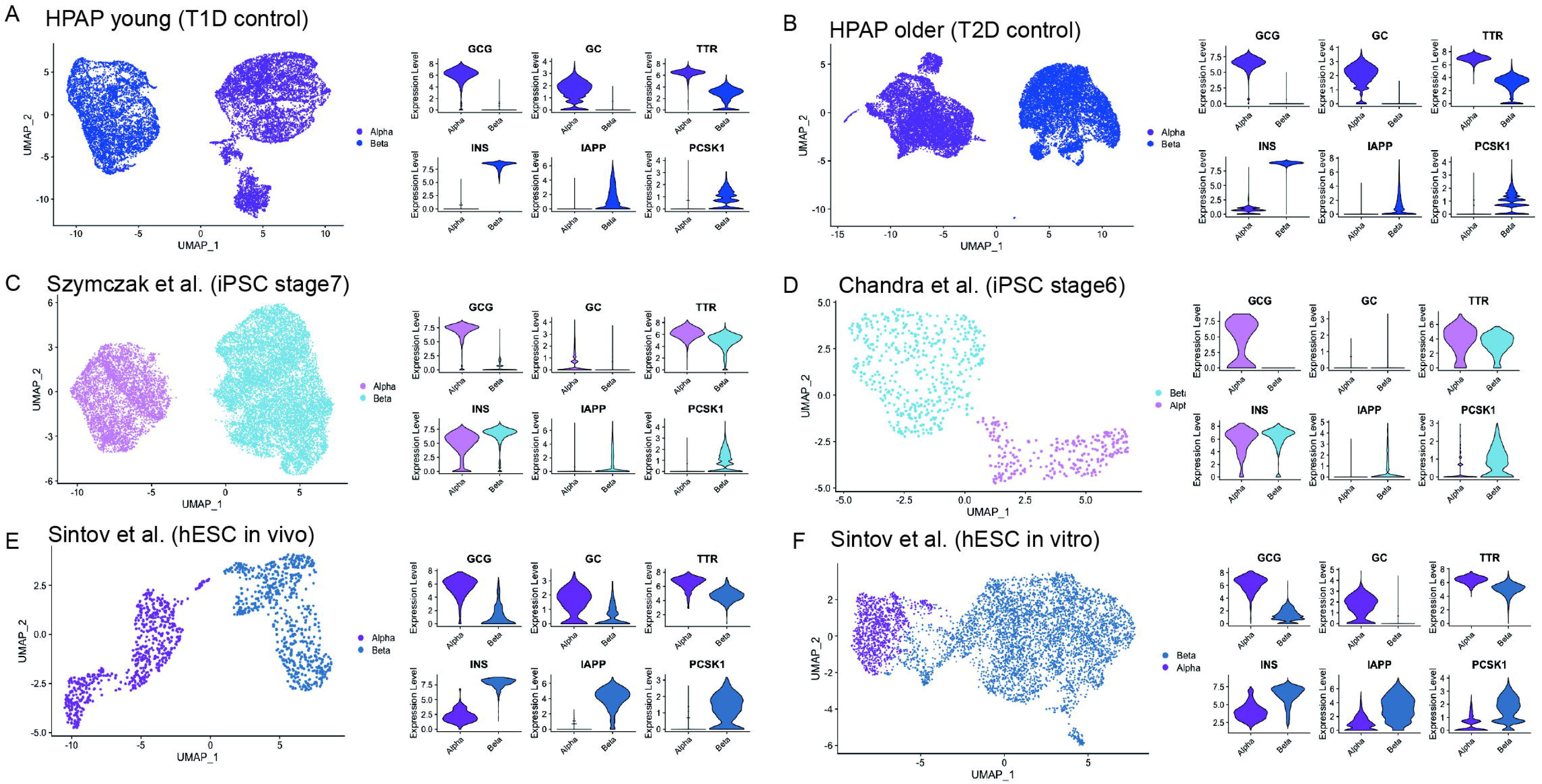
Single-cell RNA-seq profiling of basal alpha and beta cells from primary human islets and stem cell-derived islet-like cells. UMAP visualizations and violin plots showing clear separation of basal alpha- and beta cells across six single-cell RNA-seq datasets, including primary islets from HPAP donors (**A**, young T1D control; **B**, older T2D control), hiPSC-derived endocrine cells (**C**, Szymczak et al., iPSC stage 7; **D**, Chandra et al., iPSC stage 6), and hESC-derived islet-like cells (**E**, Sintov et al., hESC *in vivo*; **F**, Sintov et al., hESC *in vitro*). Violin plots display the expression of key marker genes; *GCG*, *GC* and *TTR* for alpha cells; *INS*, *IAPP* and *PCSK1* for beta cells, confirming distinct alpha- and beta-cell identities in each dataset. Only basal-state alpha and beta cells were included in this analysis. UMAP: Uniform Manifold Approximation and Projection. HPAP: Human Pancreas Analysis Program, T1D: Type 1 diabetes, T2D: Type 2 diabetes, hiPSC: human induced pluripotent stem cells, hESC: human embryonic stem cells.

### Upregulation of immune-related genes in alpha cells and *MEG3* in beta cells

We next performed differential gene expression (DGE) analysis separately for each dataset. Comparison of basal alpha and beta cells consistently revealed canonical cell identity markers among the top differentially expressed genes across the six datasets (e.g. *GCG*, *TTR*, *GC* and *ARX* for alpha cells; *INS*, *IAPP* and *PCSK1* for beta cells) (**Figure 2A-D** and **Figure S4**). The top 15 differentially expressed genes identified in the two human islet datasets had a high degree of overlap (**Figure 2A-B**). DGE analyses of alpha and beta cells from donors with one or two islet autoantibodies, as well as from donors with T1D or T2D, displayed similar patterns (**Figure S5**).

**Figure 2.**
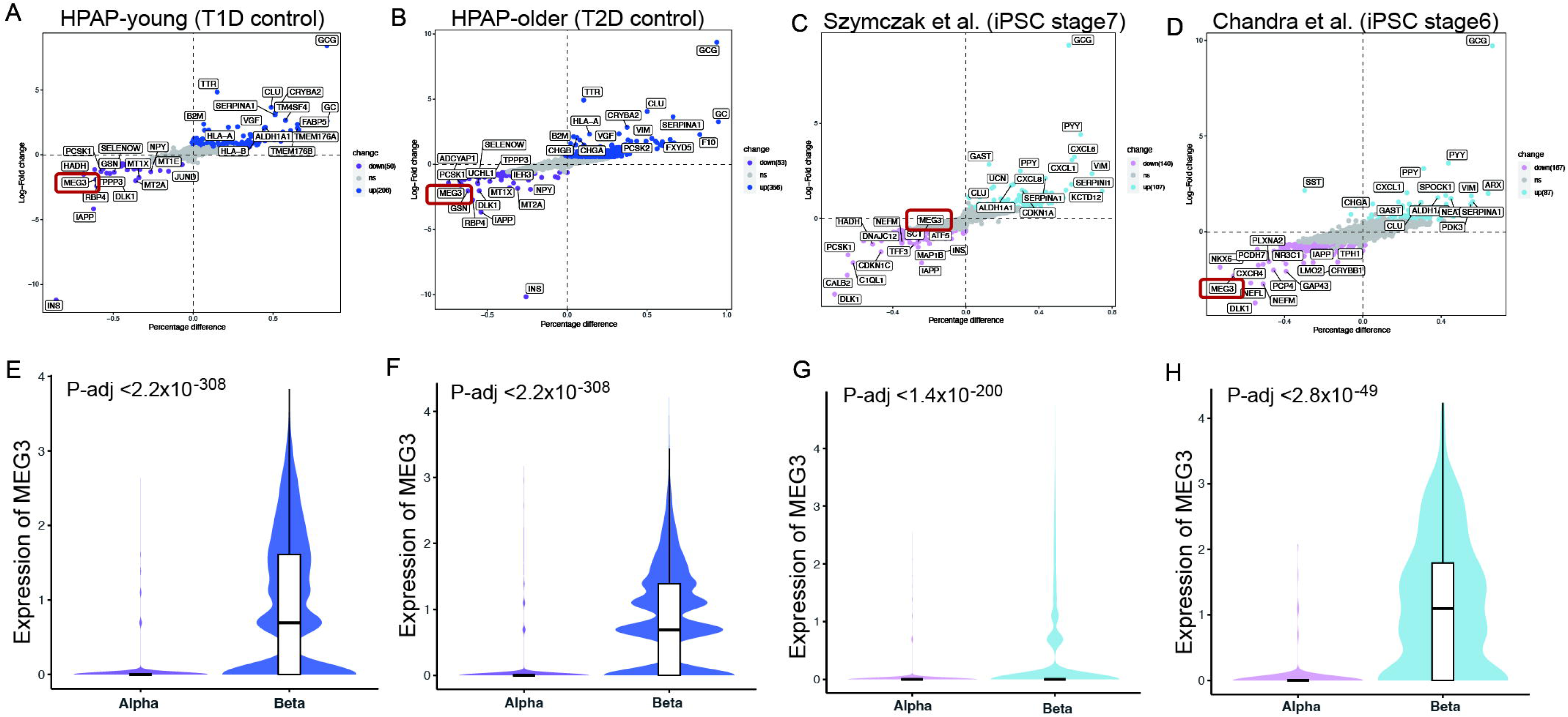
Immune-related genes are upregulated in alpha vs beta cells while *MEG3* is upregulated in beta cells compared to alpha cells from primary human islets and hiPSC-derived islet-like cells. **(A)** Differential gene expression analysis between basal alpha and beta cells obtained from HPAP-young (T1D control) donors. **(B)** Differential gene expression analysis between basal alpha and beta cells obtained from HPAP-older (T2D control) donors. **(C)** Differential gene expression analysis between basal alpha and beta cells obtained from hiPSC-derived islet-like cells (Szymczak et al., iPSC stage 7 dataset). **(D)** Differential gene expression analysis between basal alpha and beta cells obtained from hiPSC-derived islet-like cells (Chandra et al., iPSC stage 6 dataset). **(E-H)** Expression levels of *MEG3* corresponding to the datasets shown in **(A-D)**. Each dot represents a gene, plotted by log_₂_ fold change (y-axis) versus percentage difference in expression (x-axis). Genes significantly upregulated in alpha cells are shown in blue (darker blue for primary human islet samples and lighter blue for hiPSC-derived islet-like cell samples), genes downregulated in alpha cells are shown in purple (darker purple for primary human islet samples and lighter purple for hiPSC-derived islet-like cell samples), and non-significant genes are shown in grey. Selected top 15 significantly up- and downregulated genes are labelled. HPAP: Human Pancreas Analysis Program, T1D: Type 1 diabetes, T2D: Type 2 diabetes, hiPSC: human induced pluripotent stem cells.

Of interest, we found that alpha cells consistently displayed a higher “immune-like profile”, as several immune-related genes such as *HLA-A*, *HLA-B* and *B2M* were among the top 15 upregulated genes in the HPAP-young and HPAP-older datasets in both basal (**Figure 2A-B**) and disease conditions (**Figure S5**). Moreover, *JUNB*, a transcription factor regulating inflammatory cytokines expression, was strongly upregulated in hESC-derived alpha-like cells compared to beta-like cells (**Figure S4A**). In addition, the pro-apoptotic lncRNA *MEG3* was among the top upregulated genes in beta compared to alpha cells in human islets and the hiPSC-derived islet-like cells datasets under basal conditions (**Figure 2A-D**) as well as from the disease conditions (**Figure S5A, C, D**). Of note, *MEG3* expression was almost undetectable in alpha cells (**Figure 2E-H**).

### Conserved and enriched immune gene signature in alpha cells compared to beta cells

We next performed Gene Set Enrichment Analysis (GSEA) for each single-cell dataset in basal conditions. In both human islet datasets (HPAP-young and HPAP-older) and Szymczak et al. iPSC stage 7 dataset, antigen presentation-related pathways were among the top 15 enriched in alpha cells, including “Antigen presentation folding, assembly and peptide loading” and “Antigen processing cross-presentation” (**Figure 3A-C**). Additionally, interferon signaling pathways were enriched in alpha cells from the HPAP-older, Szymczak et al. iPSC stage 7 and hESC-derived islet-like cells following *in vivo* maturation (Sintov et al. hESC *in vivo*) (**Figure 3B, 3C and 3E**). Of note, “Neutrophil degranulation” was also enriched in alpha cells from hESC-derived islet-like cells differentiated *in vitro* (**Figure 3F**). These results are consistent with the strong upregulation of several immune-related genes (*HLA-A*, *HLA-B*, *B2M* and *JUNB*) in alpha cells across multiple datasets (**Figure 2, S4** and **S5**).

**Figure 3.**
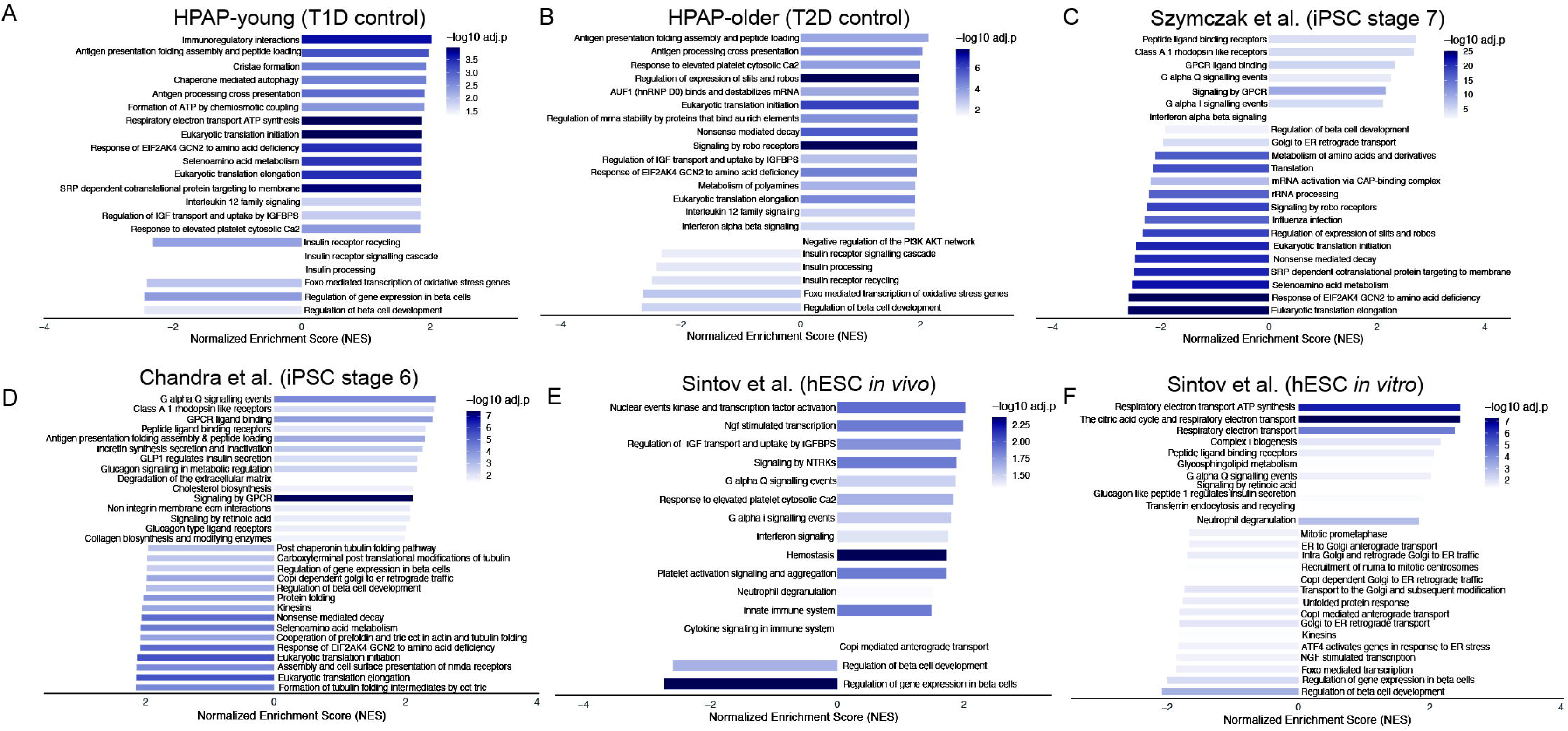
GSEA reveals conserved immune pathway enrichment in alpha cells across primary and stem cell-derived islet datasets. **(A)** GSEA of basal alpha- vs beta-cell transcriptomes from HPAP-young (T1D control) donors; **(B)** GSEA of basal alpha- vs beta-cell transcriptomes from HPAP-older (T2D control) donors; **(C)** GSEA of basal alpha- vs beta-cell transcriptomes from the dataset Szymczak et al. iPSC stage 7; **(D)** GSEA of basal alpha- vs beta-cell transcriptomes of Chandra et al. iPSC stage 6 dataset; **(E)** GSEA of basal alpha- vs beta-cell transcriptomes from the dataset Sintov et al. (hESC-derived alpha- and beta-like cells matured *in vivo*); **(F)** GSEA of basal alpha- vs beta-cell transcriptomes from the dataset Sintov et al. (hESC-derived alpha- and beta-like cells differentiated *in vitro*). Bars represent the NES, with positive values indicating alpha-cell enriched pathways and negative values indicating beta-cell enriched pathways. Color intensity reflects the significance level (– log_₁₀_ adjusted p-value). GSEA: Gene set enrichment analysis, HPAP: Human Pancreas Analysis Program, NES: normalized enrichment score.

Most of the enriched pathways in beta cells were related to beta cell function and endoplasmic reticulum (ER) activity, including “Regulation of beta cell development”, “Insulin processing”, “ER to Golgi transport”, “Eukaryotic translation”, “Protein folding” and “Unfolded protein response” (**Figure 3**). These results are expected as beta cells are equipped with a greater capacity for protein synthesis and folding machinery to support the high demand of insulin production. GSEA also revealed conserved immune pathway enrichment in alpha cells from AAB1+, AAB2+, T1D and T2D donors compared to beta cells (**Figure S6**). To further dissect the upregulation of immune-related markers in alpha versus beta cells, the expression of selected pro- and anti-inflammatory genes was analysed across alpha and beta cells from T1D, AAB2+, AAB1+ and non-diabetic donors and results showed that pro- and anti-inflammatory genes were upregulated in alpha compared to beta cells (**Figure S7A, C**). Moreover, in non-diabetic individuals both pro- and anti-inflammation gene scores were higher in alpha cells, while in T1D, beta cells showed higher pro-inflammatory scores than alpha cells (**Figure S7B, D**).

### Validation of exclusive *MEG3* expression in beta cells and enhanced immune activity in alpha cells

Single-cell transcriptomic analysis typically captures a limited number of genes per cell and may not represent the entire transcriptome (35). To further characterize the differences between alpha and beta cells, publicly available bulk RNA-seq of fluorescence-activated cell-sorted (FACS) alpha and beta cells from 12 non-diabetic donors were analyzed. Principal component analysis (PCA) confirmed clear separation between alpha and beta cells (**Figure 4A**). This was further confirmed by analyzing the expression levels of alpha- and beta-cell marker genes (**Figure S8**). DGE analysis confirmed that *MEG3* expression was significantly higher in beta cells compared to alpha cells (**Figure 4B, C**) as seen in the single-cell datasets. Alpha cells displayed enriched pathways related to mRNA splicing, including “mRNA splicing” and “Processing of capped intron-containing pre-mRNA” (**Figure 4D**). These results suggest that alpha cells may have more active post-transcriptional regulation. Meanwhile, beta cell gene expression was enriched for pathways associated with neuronal functions, including “Neuronal system”, “Neurotransmitter release cycle” and “Transmission across chemical synapses” (**Figure 4D**). These findings are consistent with the reported similarity in gene expression between pancreatic beta cells and neurons, including the shared expression of splicing regulators and splice variants (36, 37). Furthermore, alpha cells were associated with significantly higher interferon-stimulated gene (ISG) scores (9) than beta cells (**Figure 4E**).

**Figure 4.**
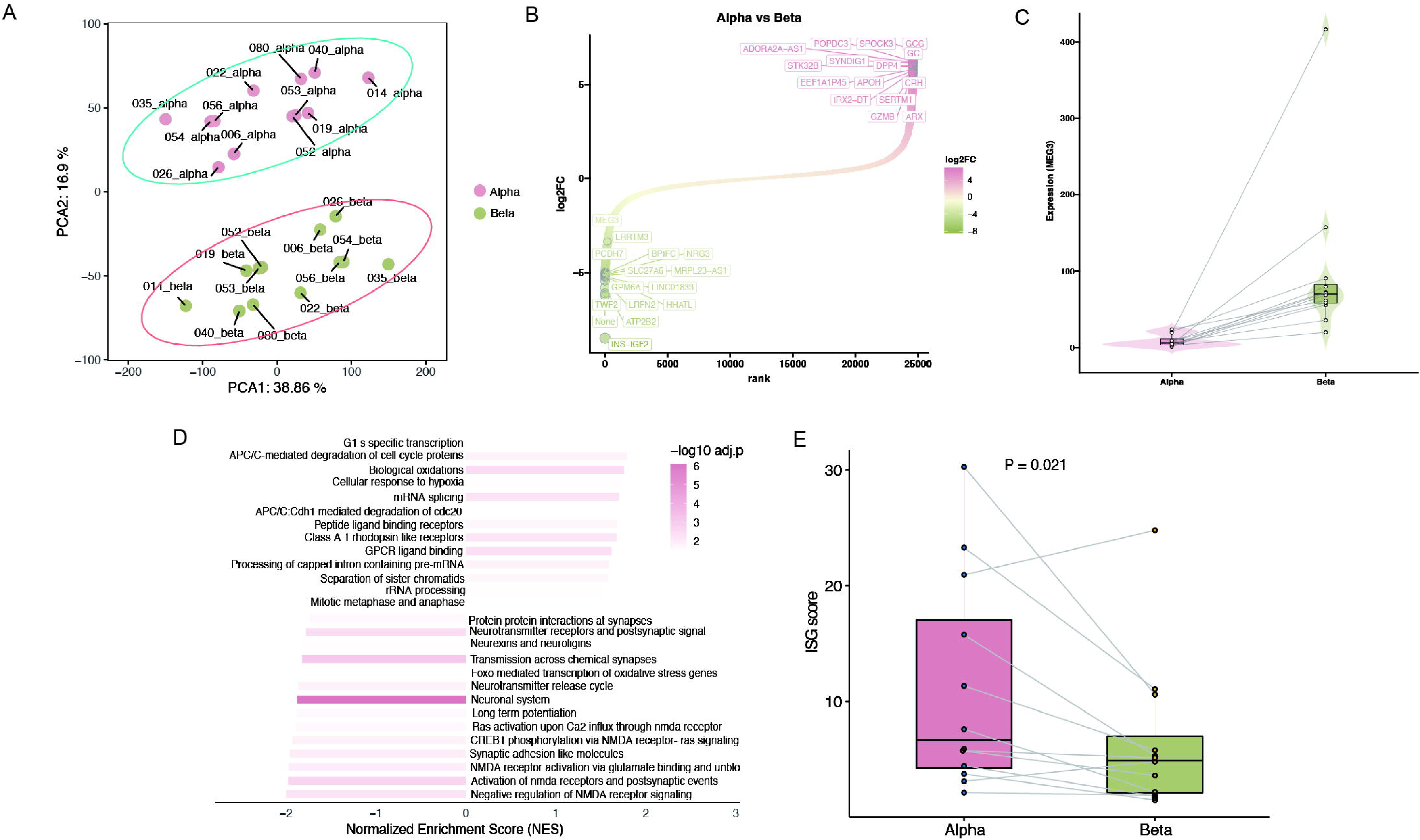
Validation of *MEG3* expression and alpha-cell enriched immune signature in FACS-sorted bulk RNA-seq data from human islets. **(A)** PCA of basal alpha and beta cells from 12 human donors using HPAP FACS-sorted bulk RNA-seq data. **(B)** Ranked gene expression differences between basal alpha and beta cells (log_₂_ fold change). **(C)** Violin plot showing *MEG3* expression levels, confirming significantly higher expression in basal beta cells compared to alpha cells. **(D)** GSEA comparing basal alpha and beta cells. **(E)** ISG scores were calculated per sample and compared using a paired two-sided t-test. PCA: Principal component analysis, HPAP: Human Pancreas Analysis Program, FACS: Fluorescence-Activated Cell Sorting, GSEA: Gene set enrichment analysis, ISG: Interferon-stimulated gene.

Together, these results corroborate our single-cell findings that *MEG3* is exclusively expressed in beta cells while alpha cells exhibit enhanced immune-related gene expression.

### Silencing *MEG3* decreases cytokine-induced apoptosis in EndoC-βH1 and hIsMTs and improves beta-cell function in cytokine-treated hIsMTs

The role of *MEG3* was further explored in human EndoC-βH1 cells exposed to a “pro-apoptotic” mix of cytokines that mimic later, adaptive immunity stages of T1D (1). The main findings were also confirmed in hIsMTs. Decreasing *MEG3* expression using two independent siRNAs decreased expression of the chemokine *CXCL10* in cytokine-exposed EndoC-βH1 cells (**Figure S9**).

Silencing *MEG3* (**Figure 5A**) in EndoC-βH1 decreased apoptosis after 48h of IFN-γ+IL-1β exposure as evaluated by 2 independent methods, namely Hoechst/Propidium iodide staining (**Figure 5B**) and Caspase 3/7 activity (**Figure 5C**). Silencing *MEG3* decreased *PDX1* levels (**Figure 5D**) but it did not significantly affect *INS* expression (**Figure 5E**) or basal insulin secretion (**Figure 5F**) from EndoC-βH1 cells. Silencing *MEG3* also did not protect EndoC-βH1 cells from the ER stressor thapsigargin (Thap)-induced apoptosis; however, untreated cells displayed decreased apoptosis levels when *MEG3* was silenced (**Figure S10A, B**). In line with the lack of protection from Thap-mediated effects, *MEG3* KD also did not affect the mRNA expression levels of the ER stress markers *BiP* and *CHOP* in these cells (**Figure S10C, D**).

**Figure 5.**
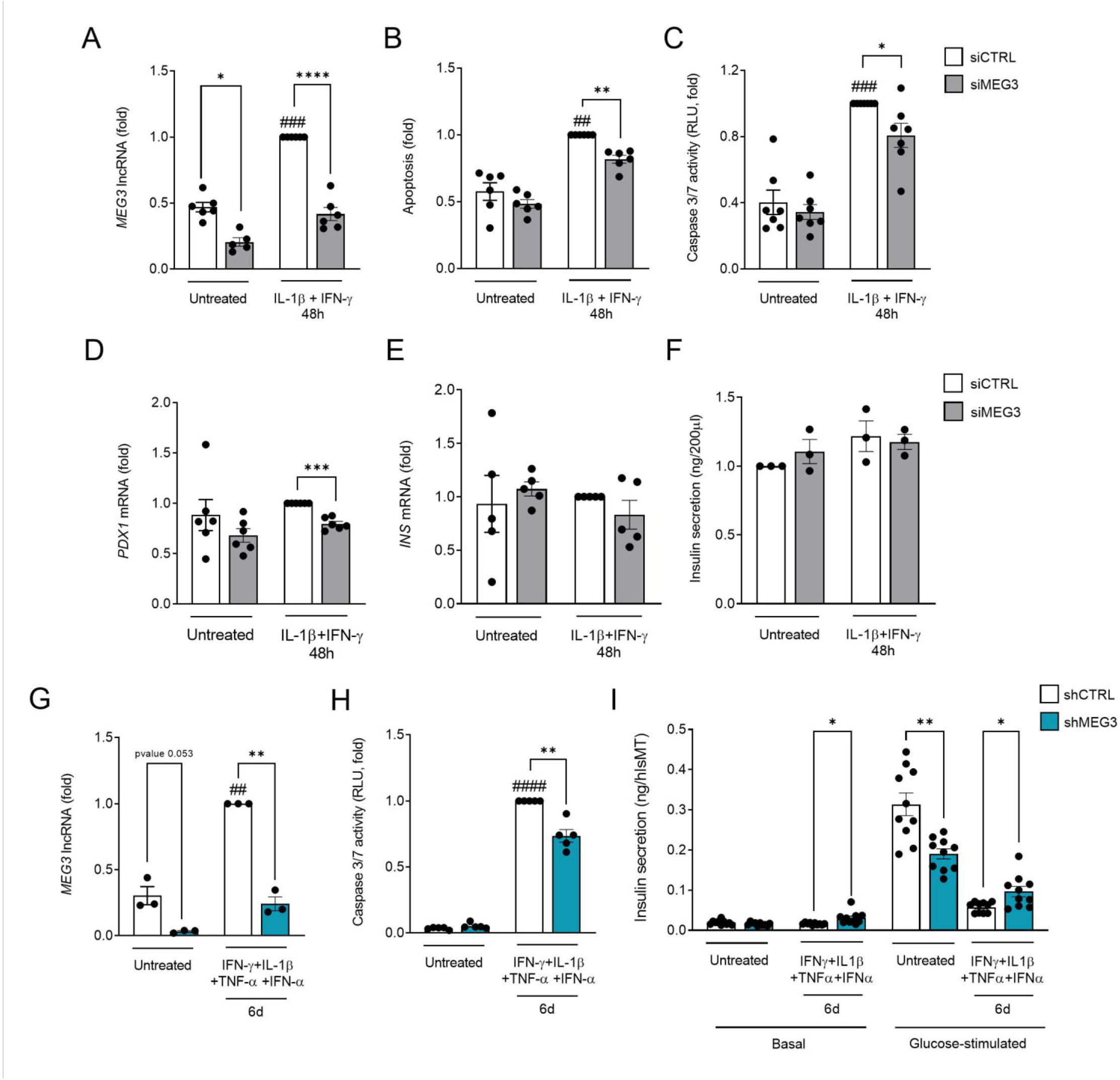
Silencing *MEG3* decreases cytokine-induced apoptosis in EndoC-βH1 and hIsMTs and improves beta-cell function in cytokine-exposed hIsMTs. **(A-F)** EndoC-βH1 cells were transfected with small interfering RNA (siCTRL or siMEG3) and **(G-I)** hIsMTs were transduced with AAVs encoding short-hairpin RNA (shCTRL or shMEG3). After recovery, cells were left untreated or exposed to IL-1β (50U/ml) +IFN-γ (1000U/ml) for 48h (for EndoC-βH1 cells) or IFN-γ (25ng/mL) +IL-1β (5ng/mL) +TNF-α (25ng/mL) +IFN-α (10ng/mL) for 6d (for hIsMTs). (A, G) *MEG3* lncRNA, (D) *PDX1* and (E) *INS* mRNA expression were assessed by RT-qPCR, normalized to the geometric mean of *ACTIN* and *VAPA* and presented as fold change compared to si/shCTRL cytokine-treated cells. **(B)** Apoptotic cells were counted using the DNA-binding dyes propidium iodide (5 mg/ml) and Hoechst 33342 (10 mg/ml). **(C, H)** Apoptosis levels were evaluated by analyzing caspase-3/7 activity. RLU are proportional to caspase-3/7 activity. **(F)** Basal and/or **(I)** glucose-stimulated insulin secretion to the supernatant was measured by ELISA. Results are means ± SEM. Each point represents an independent experiment (for EndoC-βH1 cells) or a technical replicate (for hIsMTs). *p<0.05, **p<0.01, ***p<0.001, ****p<0.0001 vs si/shCTRL, ##p<0.01, ###p<0.001, ####p<0.0001 vs untreated one-way ANOVA, Bonferroni correction. AAVs: Adeno-associated virus, HIsMTs: Human islets microtissues, RLU: Relative Luminiscence Units, SEM: Standard Error of the Mean.

*MEG3* KD using a short-hairpin RNA (shRNA) was more efficient than siRNA in hIsMTs (∼70% decrease, **Figure 5G**) and led to a reduction in cytokine-induced apoptosis as confirmed by Caspase 3/7 activity (**Figure 5H**). In addition, silencing *MEG3* increased basal and glucose-stimulated insulin secretion from cytokine-exposed hIsMTs (**Figure 5I**), demonstrating that *MEG3* KD improves islet-cell function under pro-inflammatory conditions. A limitation is that the experiments performed with hIsMTs are from a single donor and thus should be considered as indicative.

### Silencing *MEG3* decreases IFN-γ-mediated STAT1 signalling and CXCL10 expression and secretion in EndoC-βH1 cells

Western blot analyses revealed that phosphorylated STAT1 (pSTAT1) levels were reduced (albeit not significantly) after in *MEG3* KD EndoC-βH1 cells exposed to IFN-γ+IL-1β (**Figure 6A-B**). Silencing *MEG3* decreased IFN-γ-but not IFN-α-induced pSTAT1 expression (**Figure 6C**), which was confirmed by a time course including earlier time points in IFN-γ-treated EndoC-βH1 cells (**Figure 6D-F**). As a functional consequence, reduced *MEG3* levels decreased the expression and secretion of the chemokine CXCL10 (**Figure 6G, H**). These results indicate that *MEG3* plays a role in STAT1-mediated IFN-γ signalling and downstream CXCL10 expression and secretion.

**Figure 6.**
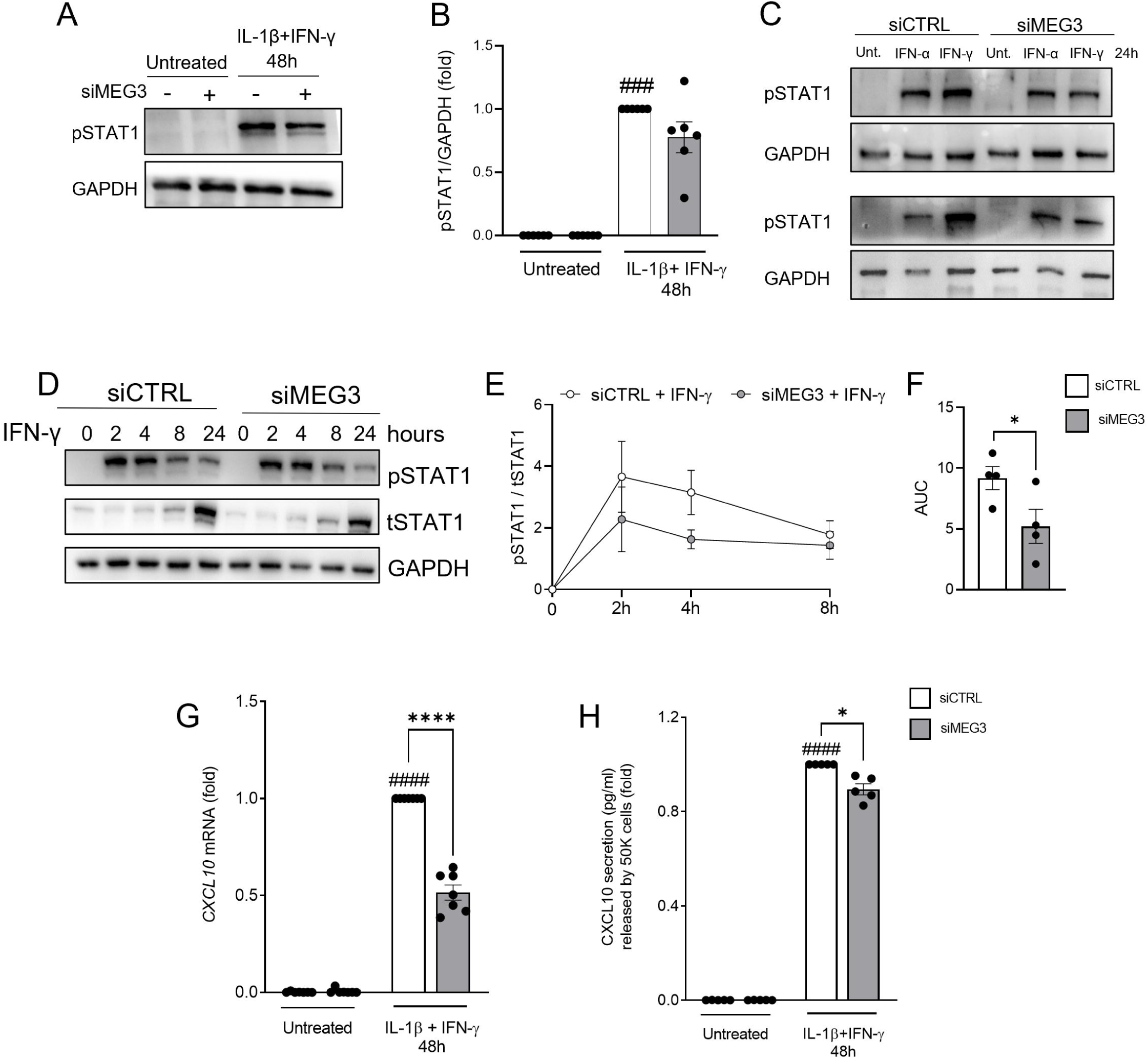
Silencing *MEG3* decreases IFN-γ-mediated STAT1 signalling and *CXCL10* expression and secretion in EndoC-βH1 cells. EndoC-βH1 cells were transfected with small interfering RNA (siCTRL or siMEG3). After recovery, cells were left untreated or exposed to either IL-1β (50U/ml) +IFN-γ (1000U/ml) for 48h, to IFN-α(1000U/ml) alone for 24h or to IFN-γ alone for 2h, 4h, 8h or 24h. **(A)** pSTAT1 and GAPDH protein expression were analyzed by immunoblot after silencing *MEG3* and exposing to the cytokines detailed in the figure. **(B)** pSTAT1 bands were quantified by densitometry and normalized to GAPDH. **(C)** Representative blots of pSTAT1 and GAPDH protein expression after silencing *MEG3* and exposing to the cytokines detailed in the figure. **(D-E)** pSTAT1, tSTAT1 and GAPDH protein expression were analyzed by immunoblot after silencing *MEG3* and exposing to IFN-γ at different time points. P-STAT1 bands were quantified by densitometry and normalized to tSTAT1. **(F)** The AUC from the figure shown in **(E)**. **(G)** *CXCL10* mRNA expression assessed by RT-qPCR, normalized to the geometric mean of *ACTIN* and *VAPA* and presented as fold change compared with siCTRL cytokine-treated cells. **(H)** CXCL10 secretion in the supernatant was measured by ELISA. Results are means ± SEM. Each point represents an independent experiment. *p<0.05, ****p<0.0001 vs siCTRL, ###p<0.001, ####p<0.0001 vs untreated one-way ANOVA, Bonferroni correction. P-STAT1: phosphorylated STAT1, tSTAT1: total STAT1, AUC: Area under the curve, SEM: Standard Error of the Mean, Unt.: untreated.

### Silencing *MEG3* decreases cytokine-induced HLA-I expression at the islet cell surface and modifies the proportion of alpha-/beta-cell fractions in hIsMTs

Cell surface expression of HLA-I is a downstream event of the STAT1 signalling pathway in beta cells. Silencing *MEG3* decreased both total and mean intensity of HLA-I surface expression in cytokine-treated hIsMTs (**Figure S11**). Additional analyses of the proportion of alpha:beta cells in hIsMTs after *MEG3* KD (**Figure S12A**) using ARX (a marker of alpha cells) and NKX6.1 (a marker of beta cells) staining indicated that silencing *MEG3* preserved the NKX6.1+ beta-cell fraction upon cytokine exposure (**Figure S12B**). Surprisingly, *MEG3* KD led to a significant decrease in ARX+/alpha cells under untreated conditions (**Figure S12C**).

## DISCUSSION

Pancreatic beta and alpha cells are hormone-secreting cells with similar developmental origins. Both are exposed to pro-inflammatory cytokines during T1D development and display similar gene expression signatures (6). However, beta cells but not alpha cells die. This difference suggests that valuable lessons for beta-cell protection could be learned from alpha cells.

Herein, we performed an integrated analysis of seven different datasets, comprising both single-cell and bulk RNA-seq. The current study compared different biological samples of alpha and beta cells from non-diabetic and diabetic donors and showed that alpha cells consistently exhibit an increased immune-like signature, with a preponderance for expression of anti-inflammatory genes (**Fig. S7**). Alpha cells also display lower expression of pro-apoptotic genes. This gene profile suggest that alpha cells are better equipped to endure immune-mediated stress compared to beta cells. In agreement with our findings, it was previously reported that rat pancreatic alpha cells are more resistant than beta cells against Coxsackievirus (CVB) infection (38). Additionally, antiviral mechanisms are up-regulated in alpha-but not in beta-like cells differentiated from hiPSCs after exposure to IFN-α (10).

Our data also revealed that T1D-associated beta cells display increased pro-inflammatory gene signatures compared to alpha cells. In addition, alpha cells express higher anti-inflammatory gene signatures under both basal and disease contexts. The concurrent activation of both pro- and anti-inflammatory pathways in alpha cells may represent an evolved adaptive mechanism that allows them to sense danger signals while preventing excessive immune activation. This feature could contribute to their increased survival in autoimmune diabetes.

Alpha cells are more resistant than beta cells to metabolic ER stress due to their higher expression of the anti-apoptotic protein BCL-xL (39). Overexpression of BCL-xL in human beta cells protected these cells against cytokine- and palmitate-induced apoptosis without hampering their function (40). Herein, we observed that the tumour suppressor *MEG3* has higher expression in beta than alpha cells across different human islets and hiPSC-derived islet-like cell datasets, from both diabetic and non-diabetic donors. Importantly, *MEG3* depletion partially protects beta/islet cells from cytokine-induced damage and apoptosis (present data). *MEG3* has been described to be downregulated in mouse models of diabetes and is linked to impairment in beta-cell identity and function (41). We presently checked the expression of beta-cell identity genes and explored beta-cell function in human cells upon *MEG3* KD. *PDX1* was significantly downregulated in cytokines-exposed EndoC-βH1 cells, but neither *INS* expression nor secretion to the culture medium were affected by *MEG3* depletion in these cells. Of note, *MEG3* KD increased basal and glucose-stimulated insulin secretion upon cytokine-exposure in hIsMTs, which may be due to reduced apoptosis and higher beta cell number as compared to shCTRL-hIsMTs exposed to cytokines. These results may be due to differences in the responses of rodent and human islets to the inflammatory environment (9, 42). Of note, *MEG3* plays a role in the activation of IFN-γ-induced phosphorylation of STAT1 (**Figure 6**). Upon IFN-γ exposure, STAT1 is phosphorylated on its tyrosine 701 residue and creates active homodimers that translocate into the nucleus and act as transcriptional activators, binding to the IFN-γ-activated sequence element at the promoter region of interferon-stimulated genes. These genes include chemokines involved in homing of immune cells to the islets in T1D such as *CXCL10* and transcription factors such as IFN regulatory factors (*IRFs*) (43). *MEG3* was previously shown to directly interact with STAT1 to regulate the promoter activity of the transcription factor IRF8 in the context of osteoclast differentiation (44). IRF8 plays an essential role in amplifying expression of IFN-γ-responsive genes initiated by STAT1 (known as “second wave of IFN-γ signalling”), an event described in immune cells (43). Whether STAT1 and *MEG3* directly interact in beta cells and whether this affects STAT1-dependent IFN-γ-directed transcriptional responses remains to be clarified. STAT1 homodimers bind to promoter elements of the transcriptional activator HLA-I transactivator, also known as *NLRC5*, and drive its transcription. *NLRC5* together with *IRF1* are key regulators of *HLA-I* expression (45, 46). The hyperexpression of HLA-I in beta cells is a hallmark of T1D and promotes neoantigen presentation and immune-mediated recognition (47, 48). Silencing *MEG3* led to a decrease in HLA-I cell-surface expression that may decrease the susceptibility of beta cells to effector T-cell attack.

ScRNA-seq has limited sensitivity compared to bulk RNA-seq, typically detecting only ∼3 000 - 5 000 highly expressed genes per cell (10, 35). This limitation motivated us to study bulk RNA-seq analysis of FACS-sorted alpha and beta cells performed at the HPAP (13), validating key findings from the single-cell analysis, including the higher *MEG3* expression in human beta cells. A limitation of our study is that we did not analyse transcript-level differences or alternative splicing (AS) regulation. We observed enriched pathways related to mRNA splicing in alpha cells, suggesting that alpha cells may exhibit more active or efficient post-transcriptional regulation. The AS landscape of thousands of genes is modulated by pro-inflammatory cytokines in human beta cells (8, 10, 49). This process may generate neoantigens, thus spreading the autoimmune response, and/or activating apoptosis- and inflammation-related proteins that contribute to beta-cell dysfunction and death (4, 50). We previously showed that the diabetes candidate gene, GLIS Family Zinc Finger 3 (*GLIS3*), modulates beta-cell apoptosis by regulating AS of the proapoptotic protein BIM (51). The *GLIS3*-regulated splicing factor *SRp55* modulates the function of the proapoptotic proteins BIM and BAX (52). The AS landscape in human alpha cells remains to be characterized and differences between alpha/beta cells in this process may open new therapeutic avenues for beta-cell protection based on splicing modulation.

In conclusion, the present findings show that combining large-scale single-cell and bulk RNA-seq analyses can identify key features underlying alpha-cell resistance to diabetes-related damage. Furthermore, the data identified the lncRNA *MEG3* as a contributing factor for beta-cell susceptibility to immune-mediated damage.

## ACKNOWLEDGMENTS

We thank A. Augenlicht, A. Musuaya, J. Capitaine, A. Bilheu, I. Millard, A.N. Belmahjoubi and N. Pachera (ULB Center for Diabetes Research, Université Libre de Bruxelles, Brussels, Belgium) for their excellent technical support, and the ULB flow cytometry platform (C. Dubois). We sincerely appreciate the HPAP Database Consortium for providing public access to the raw scRNA-seq data of human islets. The main support to D.L.E. for this project was from grants from the Breakthrough type 1 diabetes Grant Keys: 3-SRA-2023-1379-S-B and 3-IND-2024-1549-I-X. Additional support to D.L.E. was provided by the National Institutes of Health Human Islet Research Network Consortium on beta-Cell Death & Survival from Pancreatic beta-Cell Gene Networks to Therapy [HIRN-CBDS]) (grant U01 DK127786) and the National Institutes of Health Grants, NIDDK, grants RO1DK126444 and RO1DK133881-01. X.Y. was supported by the Fondation ULB, Wallonie-Bruxelles International (WBI) and the China Scholarship Council. E.M.V. was supported by the Fonds de la Recherche Scientifique (F.R.S.-FNRS, grant ID 40024397) and the Breakthrough type 1 diabetes (FY25 Fellowship ID 3-PDF-2025-1671-A-N).

## AUTHOR CONTRIBUTION STATEMENT

Conceptualization: D.L.E, X.Y. and E.M.V.; Methodology: X.Y., E.M.V., S.J.; Investigation: X.Y., E.M.V., S.J., P.Z., J.G.O., J.M.C.J., E.I., J.D.W., A.C.T., B.Y., D.L.E.; Writing of original draft: D.L.E, X.Y. and E.M.V.; Funding acquisition: D.L.E; Resources: A.C.T., B.Y.; Supervision: D.L.E. All authors have reviewed and approved the final version of the manuscript.

## ETHICS STATEMENT

Ethical approval was not required for this study.

## AVAILABILITY OF DATA AND MATERIALS

All original single-cell sequencing datasets used in this study are publicly available through the Human Pancreas Analysis Program (HPAP) portal (https://hpap.pmacs.upenn.edu/explore/download?matrix) or the Gene Expression Omnibus (GEO) under accession numbers: <u>GSE203384</u>, <u>GSE190726</u>, <u>GSE200083</u>, and <u>GSE200084</u>. The original bulk RNA sequencing reads of FACS-sorted basal alpha and beta cells was obtained from HPAP (https://hpap.pmacs.upenn.edu/explore/download?matrix) (2024 release).

