## Supplemental Figures for "Differential immune- and apoptosis-related gene signatures in pancreatic alpha and beta cells contribute to their fate in type 1 diabetes"

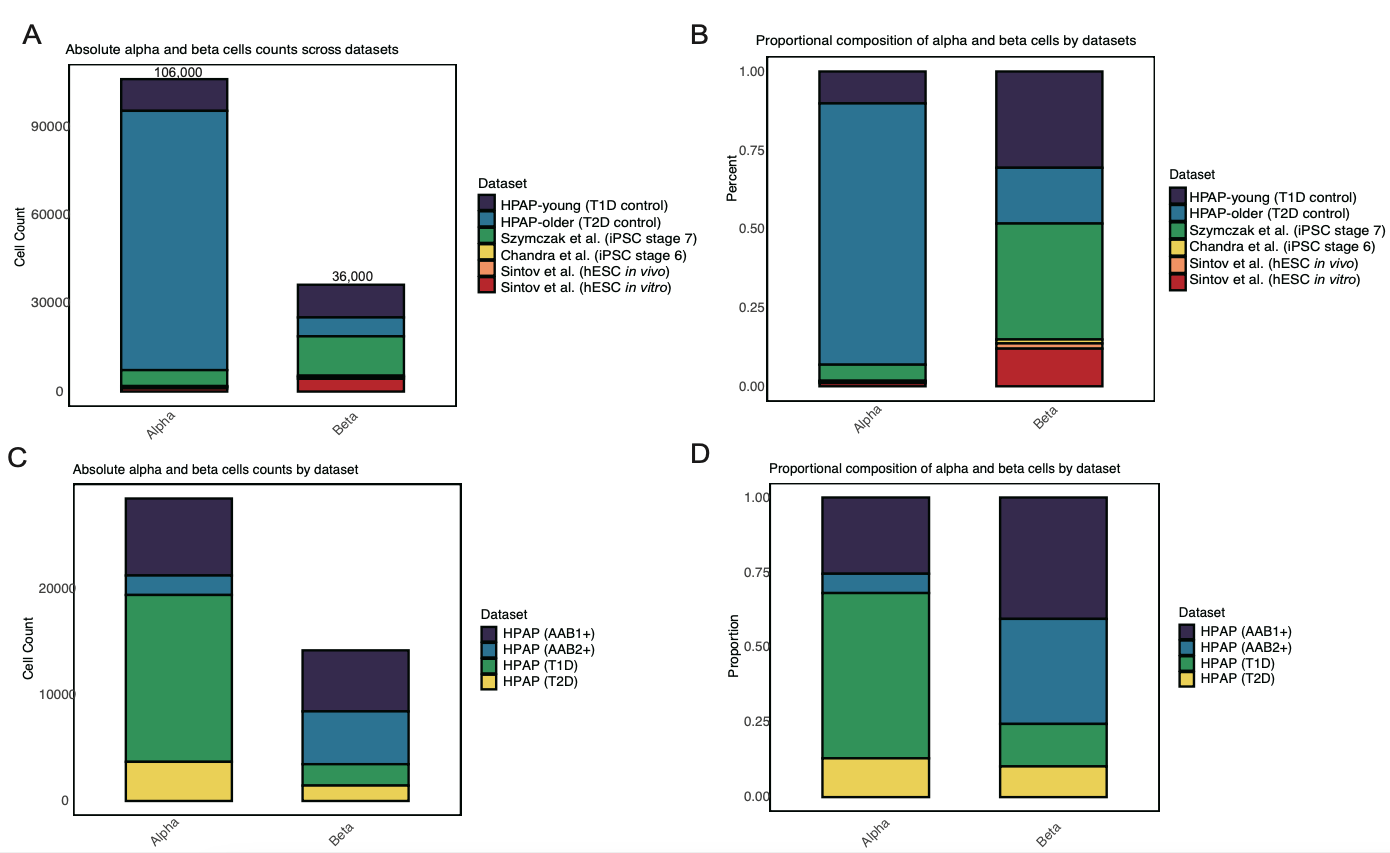

**Figure S1. Counts and relative composition of alpha and beta cells across primary human islets and stem cell-derived islet-like cell datasets. (A)** Absolute counts of basal alpha and beta cells obtained from six single-cell RNA-seq datasets, including primary human islets (HPAP donors: young T1D control and older T2D control), hiPSC-derived islet-like cells (Szymczak et al., iPSC stage 7; Chandra et al., iPSC stage 6), and hESC-derived cells (Sintov et al., hESC *in vivo* and *in vitro*). **(B)** Normalized proportions of alpha and beta cells across datasets, showing relative cellular composition within each lineage. **(C)** Absolute counts of alpha and beta cells from AAB1+, AAB2+, T1D and T2D donors. **(D)** Relative composition of alpha and beta cells from AAB1+, AAB2+, T1D and T2D donors. T1D: Type 1 diabetes, T2D: Type 2 diabetes, hiPSC: human induced pluripotent stem cells, hESC: human embryonic stem cells. AAB1+: donors positive for one islet autoantibody. AAB2+: donors positive for two or more islet autoantibodies.

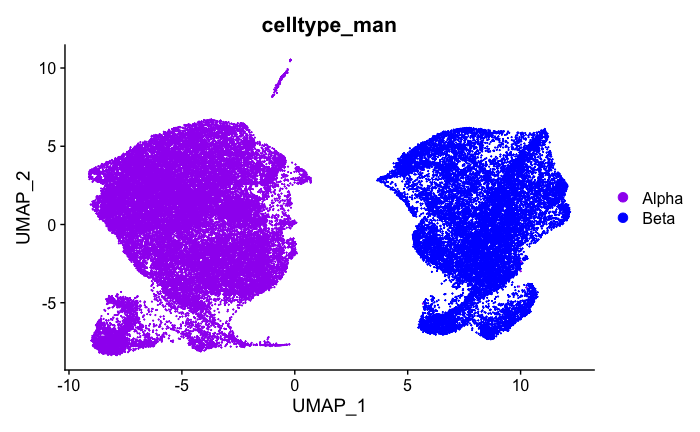

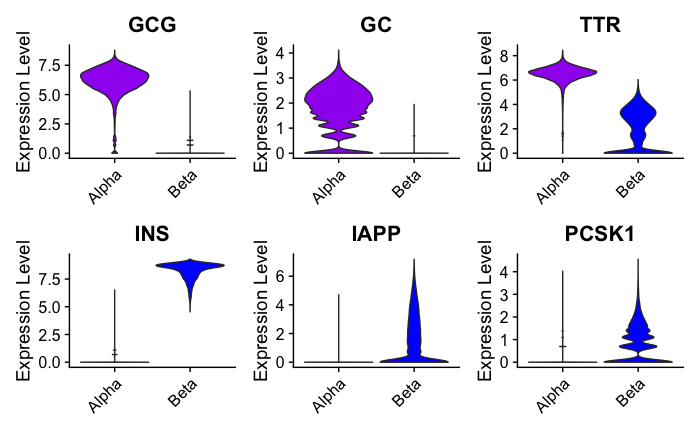

A

B

HPAP T1D, AAB1+, AAB2+, ND

HPAP T2D, ND

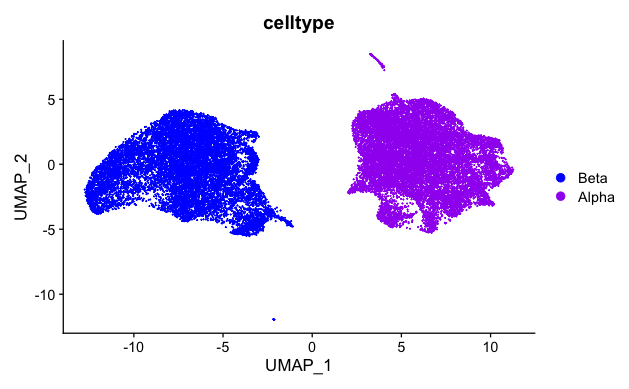

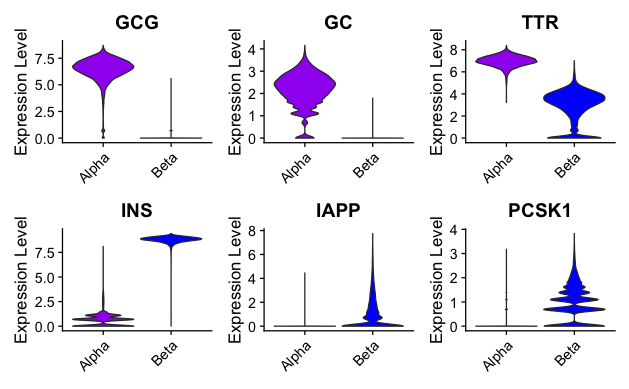

**Figure S2. Single-cell RNA-seq profiling of alpha and beta cells from primary human islets.** UMAP visualizations and violin plots showing clear separation of alpha and beta cells, including primary islets from HPAP donors: **(A)** T1D donors, donors with two or one islet autoantibodies and control donors; **(B)** T2D and control donors. Violin plots display the expression of key marker genes: *GCG*, *GC* and *TTR* for alpha cells; *INS*, *IAPP* and *PCSK1* for beta cells, confirming distinct alpha- and beta-cell identities in each dataset. UMAP: Uniform Manifold Approximation and Projection. HPAP: Human Pancreas Analysis Program, T1D: Type 1 diabetes, T2D: Type 2 diabetes. AAB1+: donors positive for one islet autoantibody. AAB2+: donors positive for two or more islet autoantibodies, ND: non-diabetic donors.

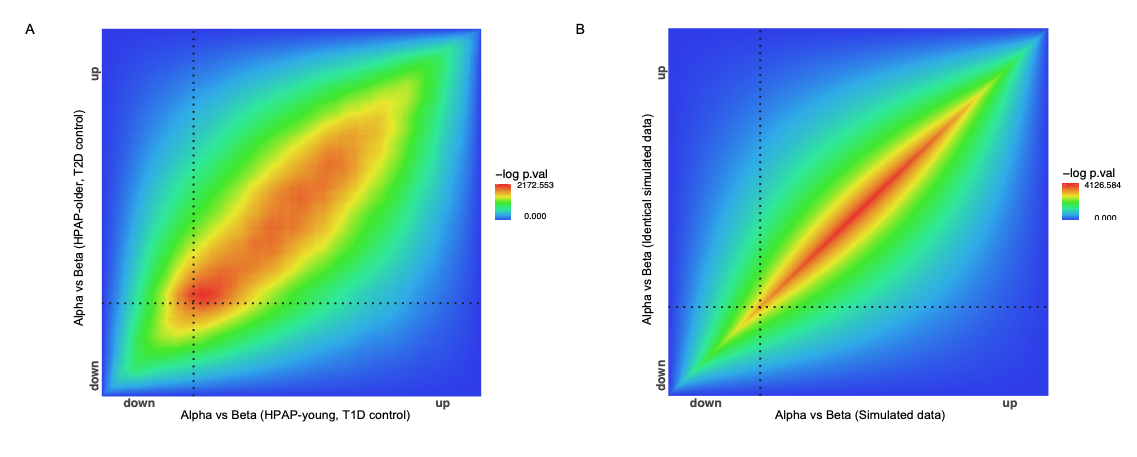

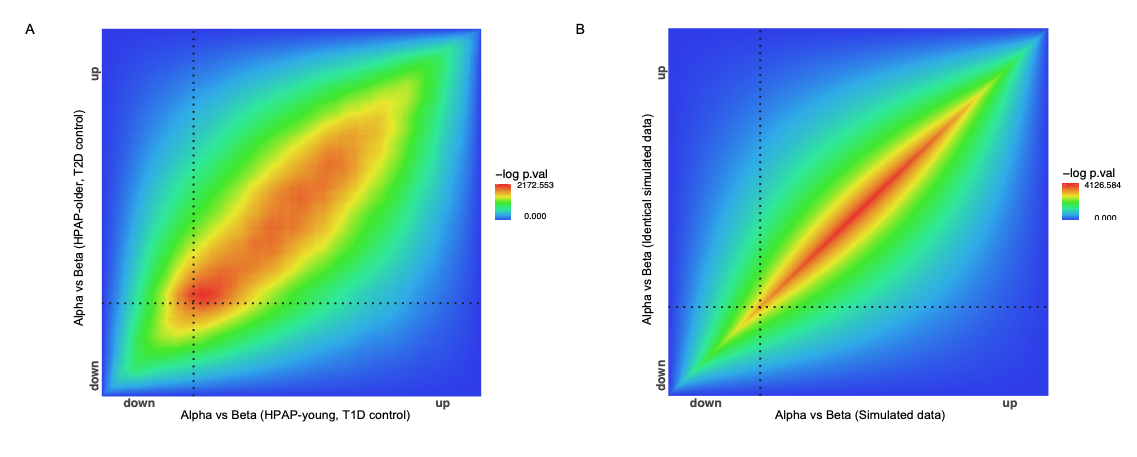

**Figure S3. Comparative analysis of basal alpha- vs beta-cell transcriptional differences between young and older donors. (A)** Two-dimensional RRHO-RedRibbon level map of basal alpha vs beta cells between HPAP-young (T1D control) and HPAP-older donors (T2D control). **(B)** Reference correlation using simulated data with identical fold-change distributions, representing a theoretical perfect correlation scenario. Colours represent the -log(p-value) for RRHO-RedRibbon analysis, from low (blue) to high (red). HPAP: Human Pancreas Analysis Program, RRHO: Rank-rank Hypergeometric Overlap, T1D: Type 1 diabetes, T2D: Type 2 diabetes.

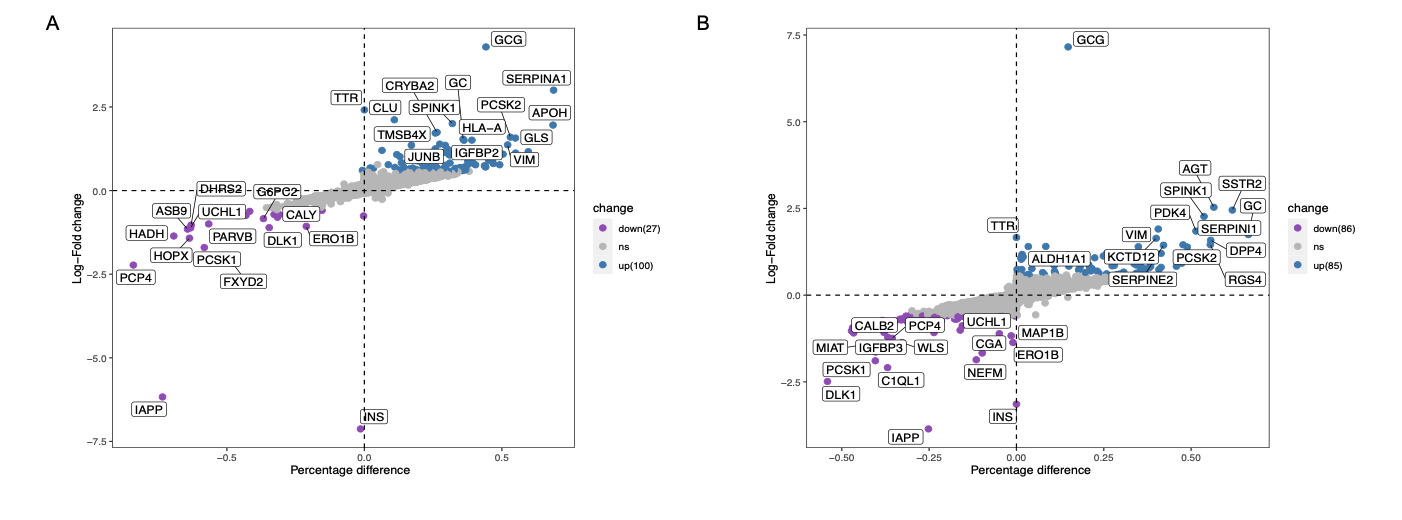

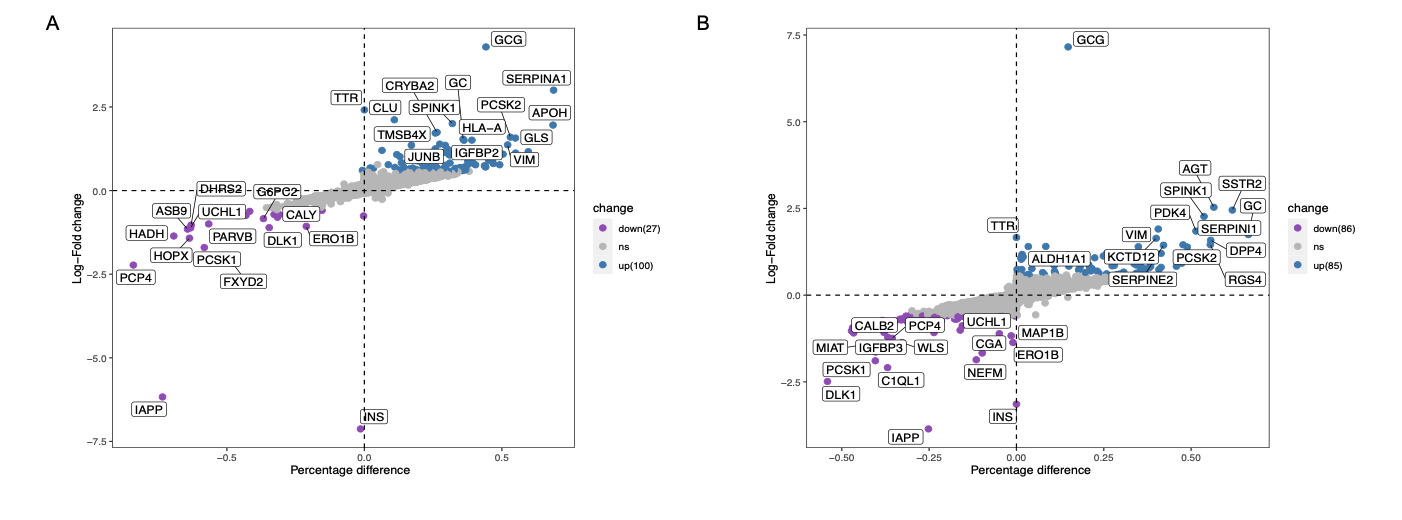

Sintov et al. (hESC *in vitro*)

Sintov et al. (hESC *in vivo*)

**Figure S4. Differential gene expression analysis between basal alpha and beta cells derived from hESCs. (A)** Basal alpha and beta cells were obtained from single-cell RNA-seq data of *in vivo* matured hESC-derived islet-like cells isolated from humanized mouse after transplantation (Sintov et al. hESC in vivo). **(B)** Basal alpha and beta cells were obtained from single-cell RNA-seq data of *in vitro* differentiated hESC-derived islet-like cells (Sintov et al. hESC in vitro). Each dot represents a gene, plotted by log₂ fold change (y-axis) vs percentage difference in expression (x-axis). Genes significantly upregulated in alpha cells are shown in blue, genes downregulated in alpha cells are shown in purple and non-significant genes are shown in grey. Selected top 15 significantly up- and downregulated genes are labelled. hESC: human embryonic stem cells.

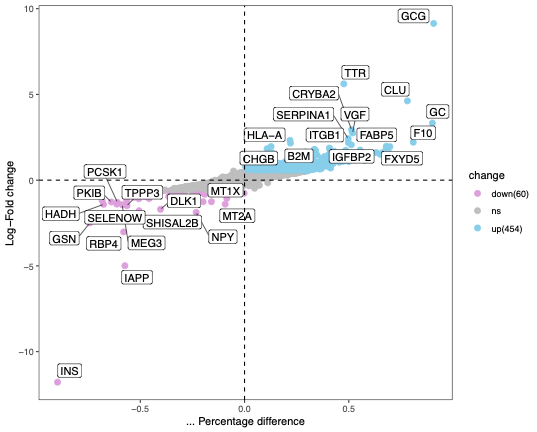

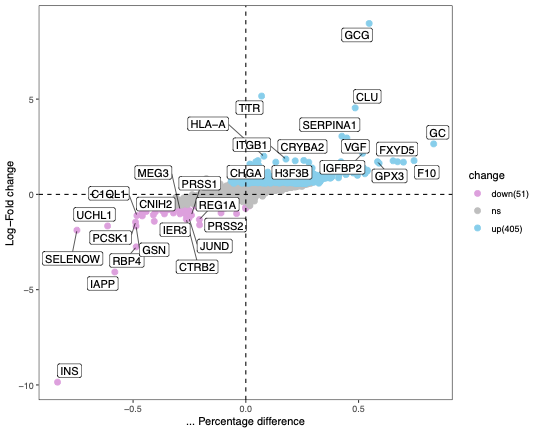

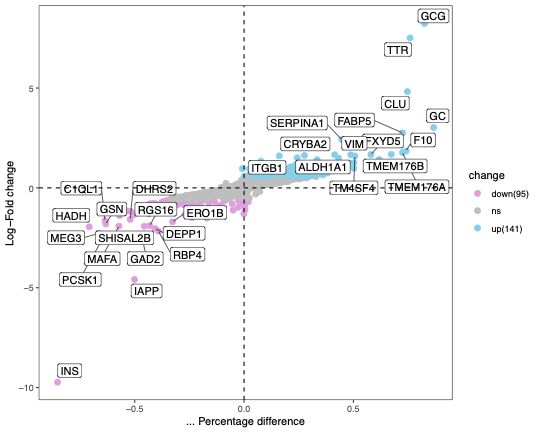

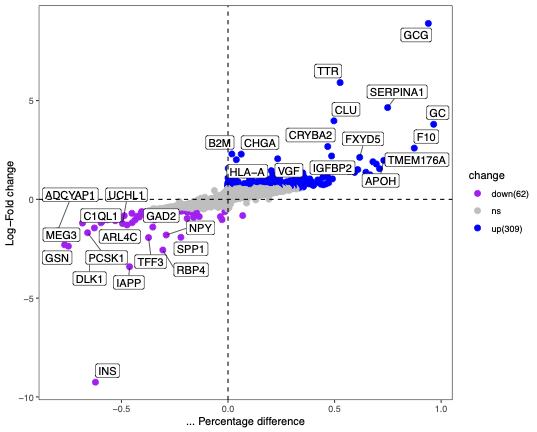

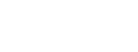

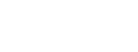

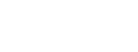

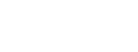

A

B

C

D

AAB2+ donors

AAB1+ donors

T2D donors

T1D donors

**Figure S5. Differential gene expression analysis between alpha and beta cells from AAB1+, AAB2+, T1D, and T2D human islet donors.** Alpha and beta cells from AAB1+ **(A)**, AAB2+ **(B)**, T1D **(C)** and T2D **(D)** donors were compared. Each dot represents a gene, plotted by log₂ fold change (y-axis) versus percentage difference in expression (x-axis). Genes significantly upregulated in alpha cells from the HPAP T1D cohort are shown in sky blue, and those from the HPAP T2D cohort are shown in blue. Downregulated genes are shown in purple (T1D) and plum (T2D), while non-significant genes are shown in grey. The top 15 significantly up- and downregulated genes are labelled. HPAP: Human Pancreas Analysis Program, T1D: Type 1 diabetes, T2D: Type 2 diabetes, AAB1+: donors positive for one islet autoantibody, AAB2+: donors positive for two or more islet autoantibodies.

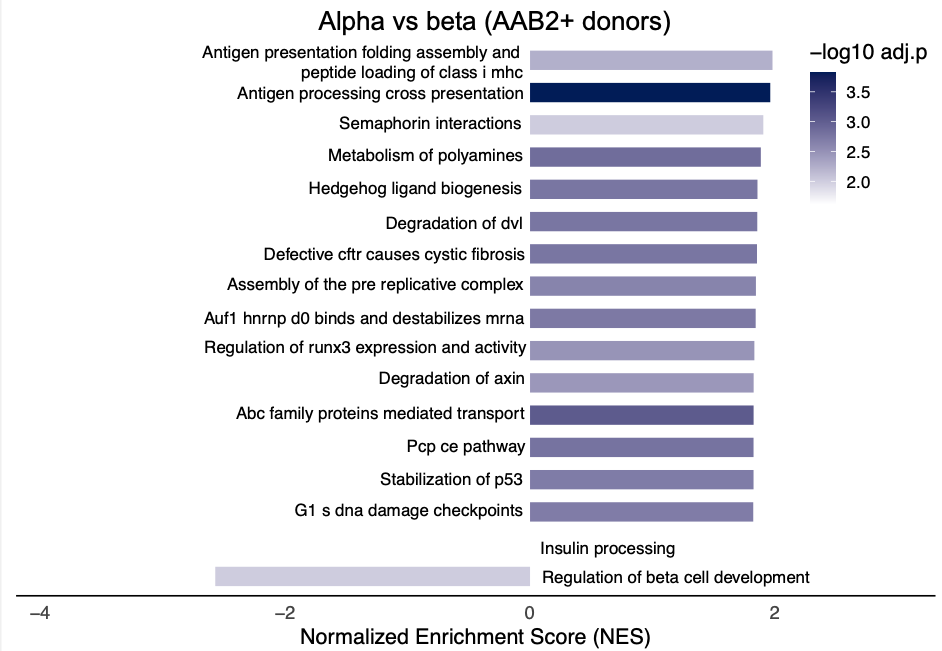

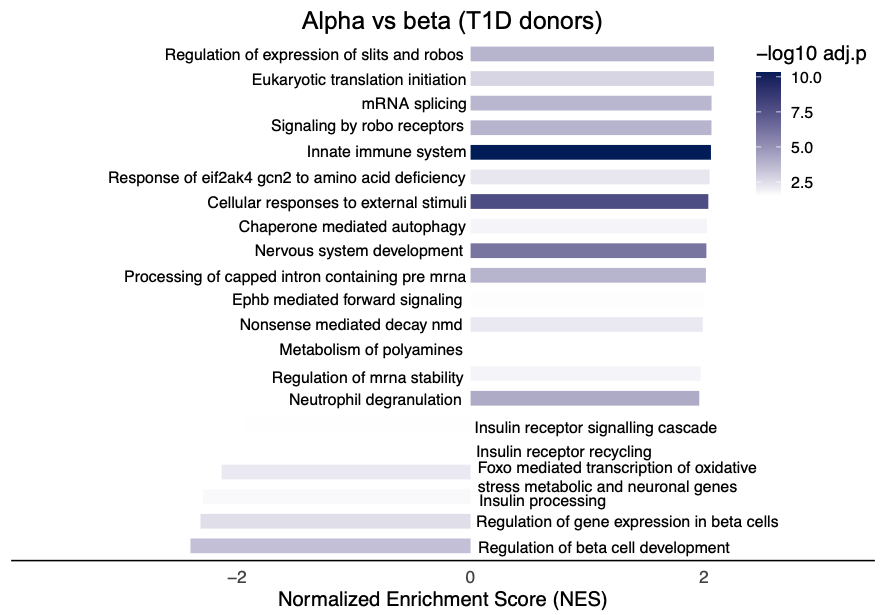

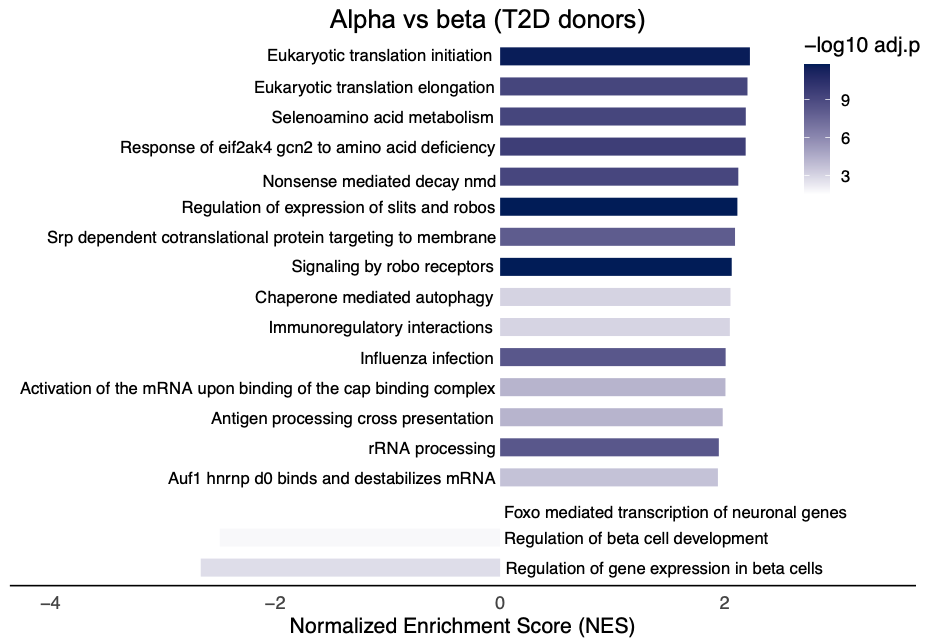

A

B

C

D

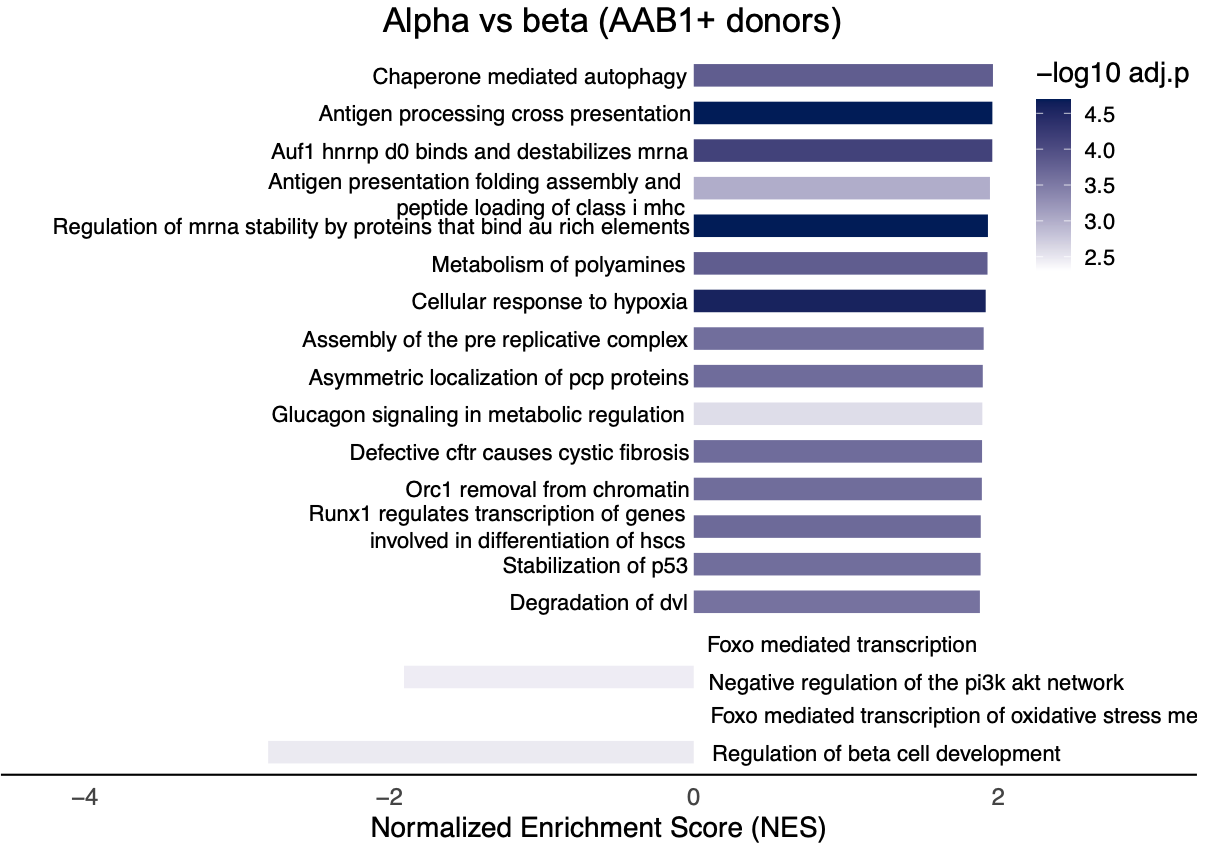

C

D

**Figure S6. GSEA reveals conserved immune pathway enrichment in alpha cells from AAB1+, AAB2+, T1D and T2D donors compared to beta cells. (A)** GSEA of AAB1+ alpha- vs beta-cell transcriptomes; **(B)** GSEA of AAB2+ alpha- vs beta-cell transcriptomes; **(C)** GSEA of T1D alpha- vs beta-cell transcriptomes; **(D)** GSEA of T2D alpha- vs beta-cell transcriptomes. All donors are from the HPAP cohort. Bars represent the NES, with positive values indicating alpha-cell enriched pathways and negative values indicating beta-cell enriched pathways. Color intensity reflects the significance level (-log₁₀ adjusted p-value). GSEA: Gene set enrichment analysis, HPAP: Human Pancreas Analysis Program, NES: normalized enrichment score. AAB1+: donors positive for one islet autoantibody. AAB2+: donors positive for two or more islet autoantibodies.

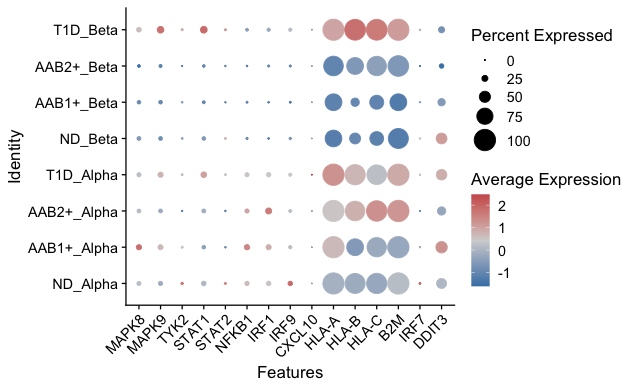

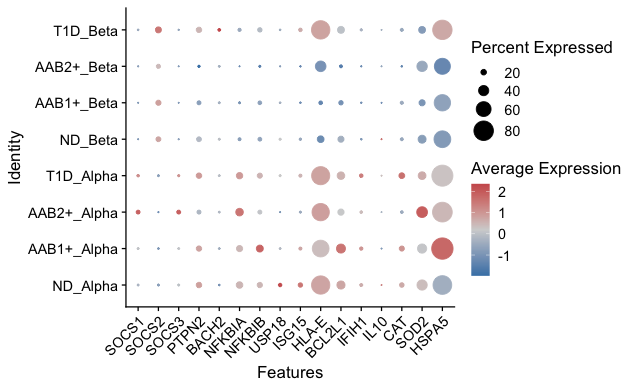

Pro-inflammation

Anti-inflammation

A

C

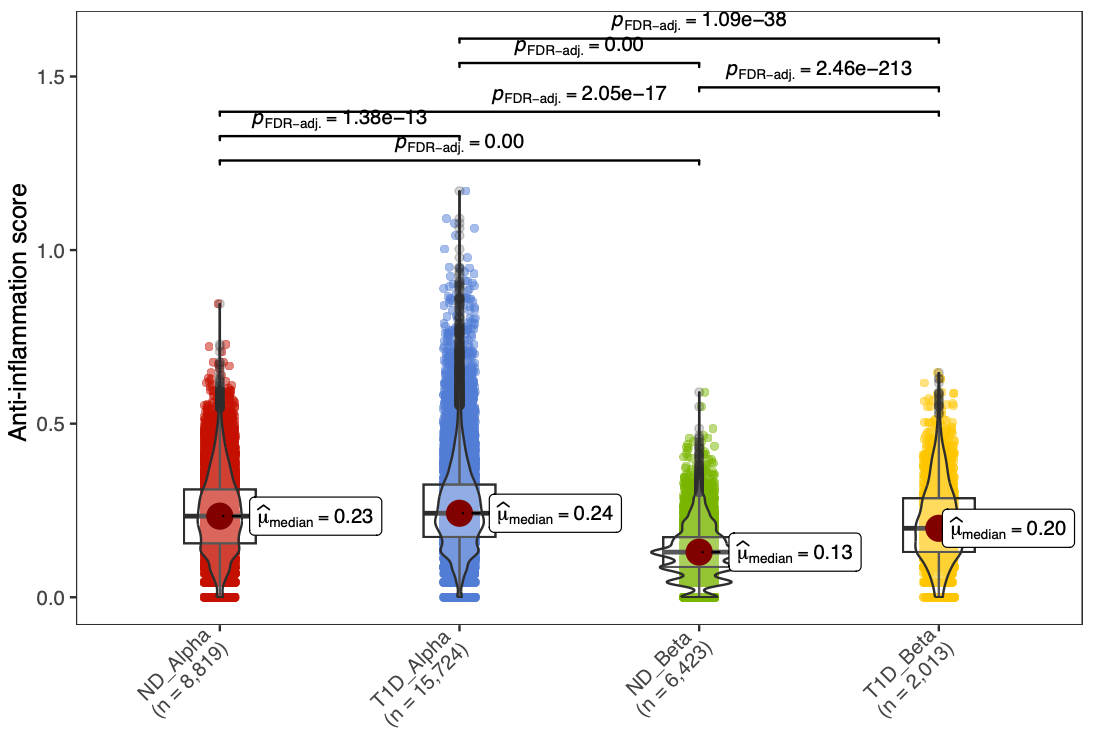

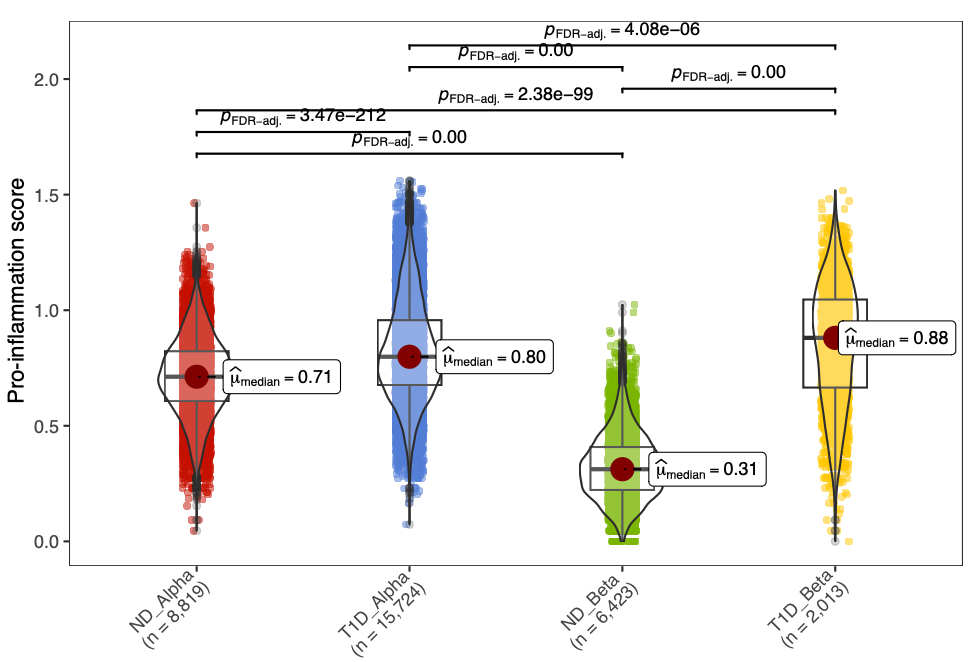

B

D

**Figure S7. Expression of pro- and anti-inflammatory genes in alpha and beta cells from HPAP donors with T1D, AAB2+, AAB1+ and controls.** **(A)** Dot plot showing the expression of selected pro-inflammatory genes across alpha and beta cells from T1D, AAB2+, AAB1+, and ND donors. Dot size represents the percentage of cells expressing each gene, and colour indicates average expression (scaled). **(B)** Violin plot displaying the pro-inflammation gene score per group, with median and interquartile range shown. Statistical comparisons were performed using two-sided Wilcoxon tests. **(C)** Dot plot showing the expression of selected anti-inflammatory genes across the same donor groups. **(D)** Violin plot showing the anti-inflammation gene score per group, with statistical comparisons as in **(B)**. T1D: Type 1 Diabetes, AAB1+: donors positive for one islet autoantibody, AAB2+: donors positive for two or more islet autoantibodies, ND: non-diabetic controls, HPAP: Human Pancreas Analysis Program.

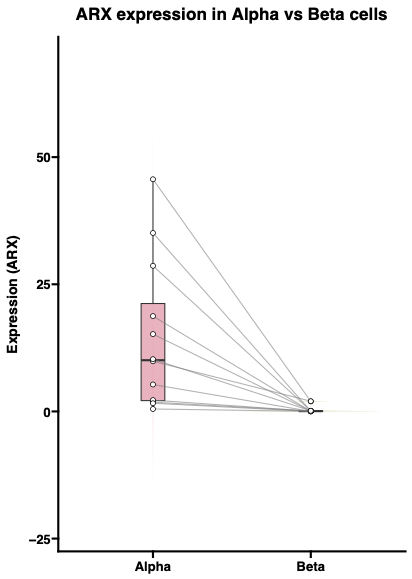

A

TPM for ARX

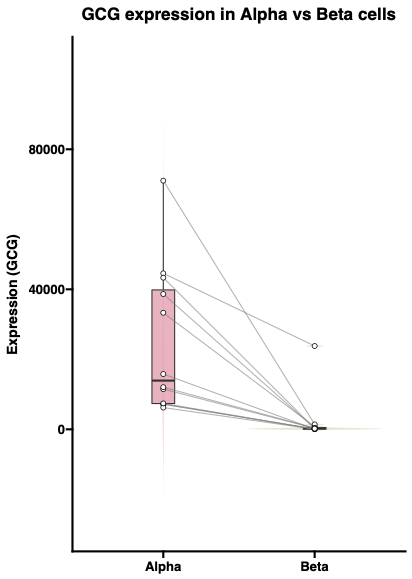

B

P-adj: 3.9x10^-21^

TPM for GCG

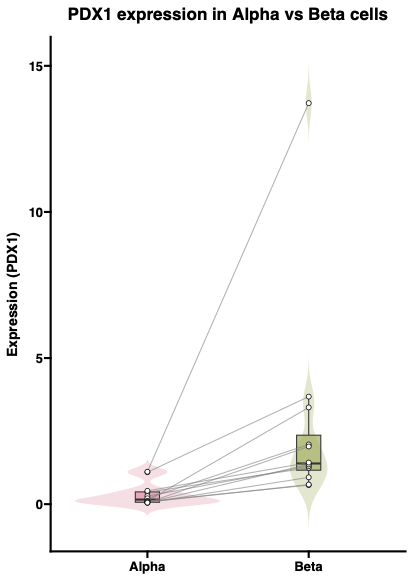

C

TPM for PDX1

D

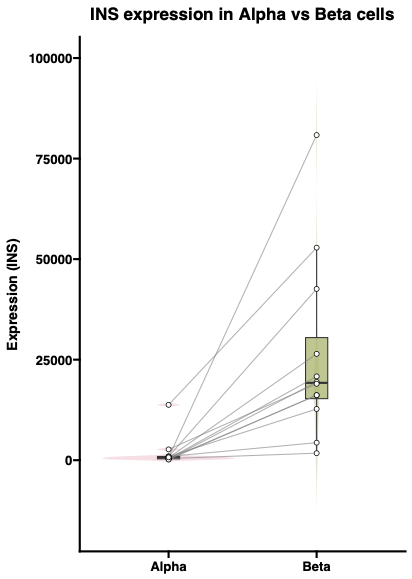

TPM for INS

P-adj: 1.7x10^-31^

P-adj: 1.2x10^-25^

P-adj: 2.9x10^-8^

**Figure S8.** **Expression levels of alpha- and beta-cell marker genes in HPAP FACS-sorted alpha and beta cells.** Boxplots showing TPM values for alpha-cell markers (*ARX* and *GCG*) and beta-cell markers (*PDX1* and *INS*) across 12 donors. Bulk RNA-seq data were obtained from FACS-sorted basal alpha and beta cells from the HPAP. Paired data points for each donor are connected by lines, indicating that the alpha and beta cells are derived from the same donor. Statistical analysis was performed using Wald tests from DESeq2, and P-values were adjusted using the Benjamini-Hochberg method. HPAP: Human Pancreas Analysis Program, FACS: Fluorescence-activated cell sorting, TPM: transcripts per million.

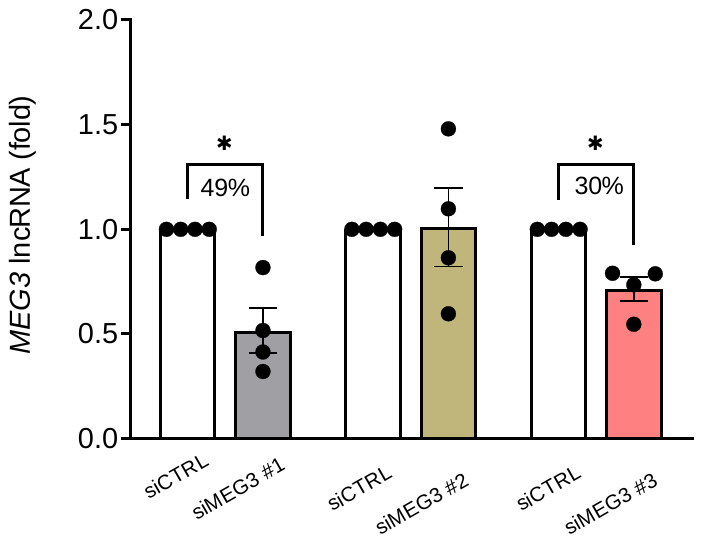

A

B

C

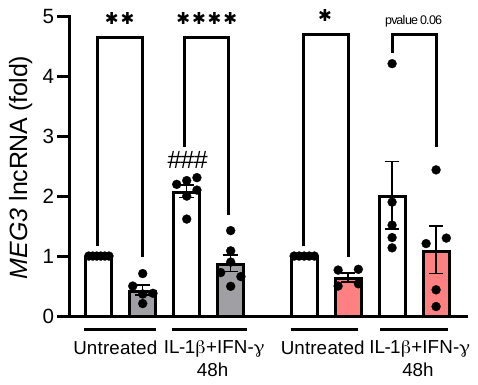

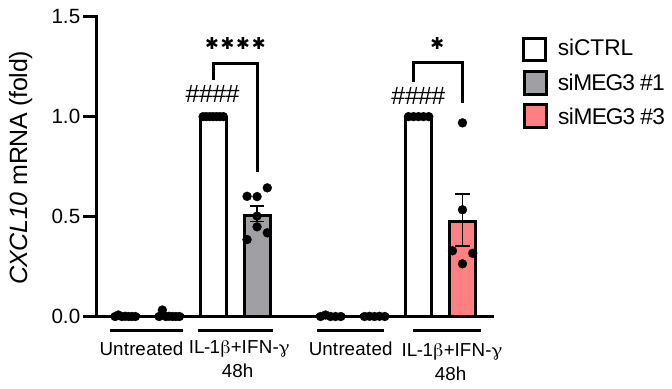

**Figure S9. Validation of *MEG3* silencing. (A)** EndoC-βH1 cells were transfected with small interfering RNA (siCTRL, siMEG3 #1, siMEG3 #2 or siMEG3 #3). After 24h of recovery, cells were left untreated for 48h. *MEG3* lncRNA expression was assessed by RT-qPCR, normalized to the geometric mean of *ACTIN* and *VAPA* and presented as fold change compared with siCTRL cells. SiMEG3 #1, the most efficient siRNA, was used in all subsequent experiment. **(B, C)** EndoC-βH1 cells were transfected with small interfering RNA (siCTRL, siMEG3 #1 or siMEG3 #3). After 24h of recovery, cells were left untreated or exposed to IL-1β (50U/ml) +IFN-γ (1000U/ml) for 48h. **(B)** *MEG3* lncRNA and **(C)** *CXCL10* mRNA expression was assessed by RT-qPCR, normalized to the geometric mean of *ACTIN* and *VAPA* and presented as fold change compared with siCTRL cells untreated or treated with cytokines. Results are means ± SEM. Each point represents an independent experiment. *p<0.05, **p<0.01, ****p<0.0001 vs siCTRL, ###p<0.001, ####p<0.0001 vs untreated one-way ANOVA, Bonferroni correction. SEM: Standard Error of the Mean.

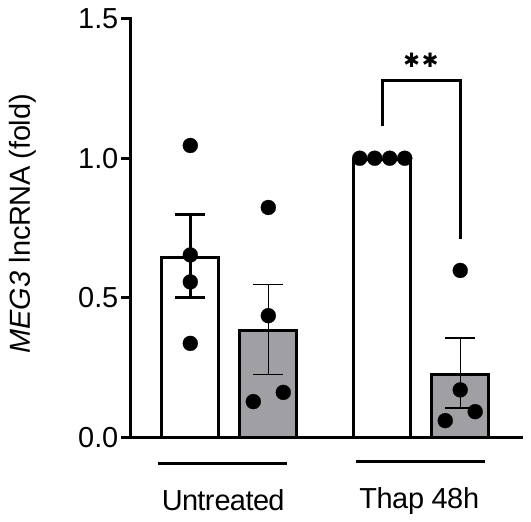

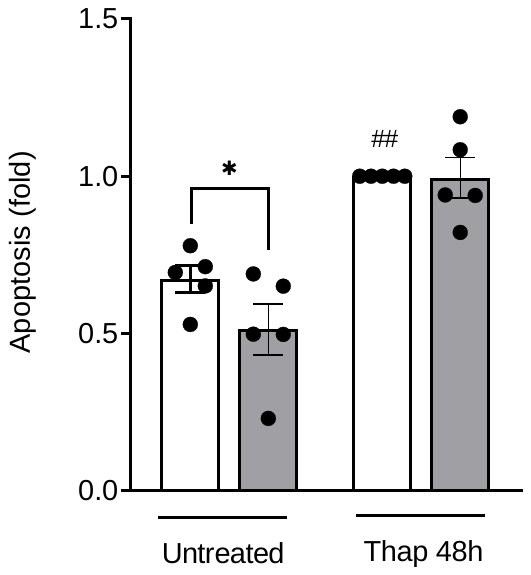

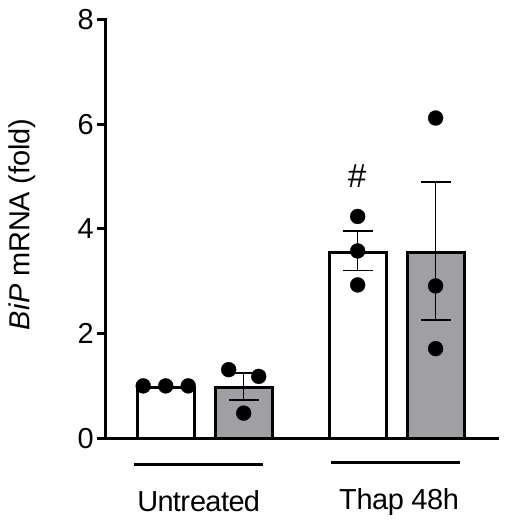

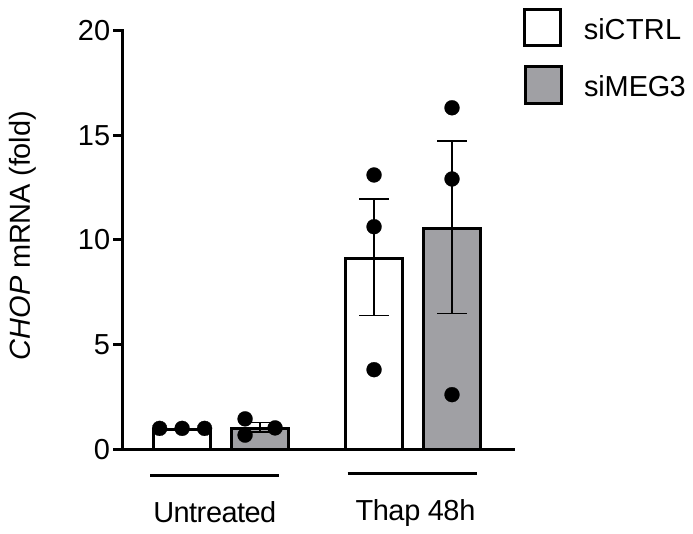

A

B

C

D

**Figure S10. Silencing of *MEG3* does not protect EndoC-βH1 cells from thapsigargin (Thap)-induced gene expression nor apoptosis.** EndoC-βH1 cells were transfected with small interfering RNA (siCTRL or siMEG3). After 24h of recovery, cells were left untreated or exposed to the ER stressor Thap (1μM, 48h). **(A)** *MEG3* lncRNA, **(C)** *BiP* mRNA and **(D)** *CHOP* mRNA expression was assessed by RT-qPCR, normalized to the geometric mean of *ACTIN* and *VAPA* and presented as fold change compared with siCTRL untreated or Thap-treated cells. **(B)** Apoptotic cells were counted with the DNA-binding dyes propidium iodide (5 mg/ml) and Hoechst 33342 (10 mg/ml). Results are means ± SEM. Each point represents an independent experiment. *p<0.05, **p<0.01, vs siCTRL, #p<0.05, ##p<0.01 vs untreated, one-way ANOVA, Bonferroni correction. Thap: thapsigargin, SEM: Standard Error of the Mean.

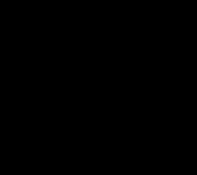

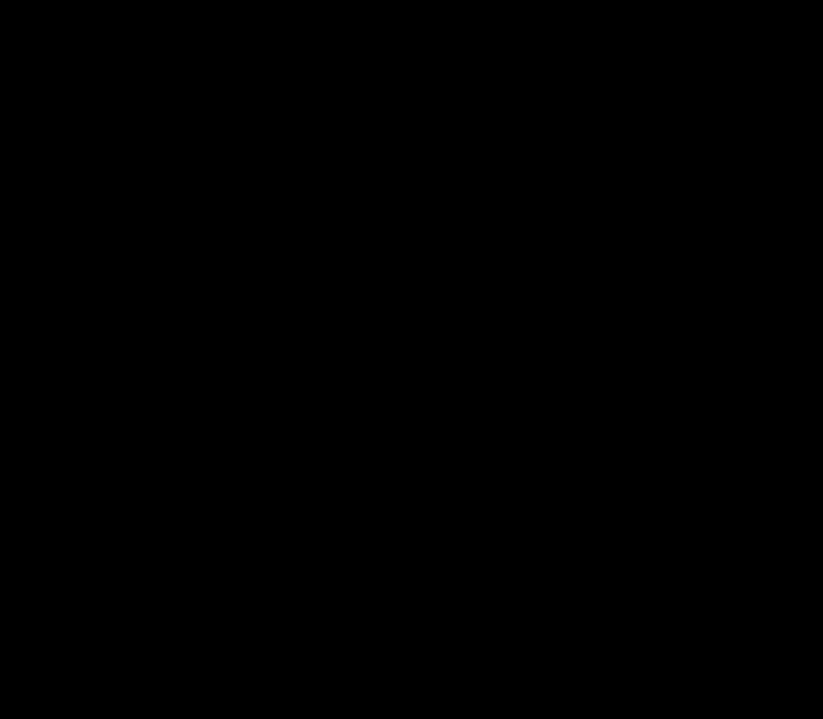

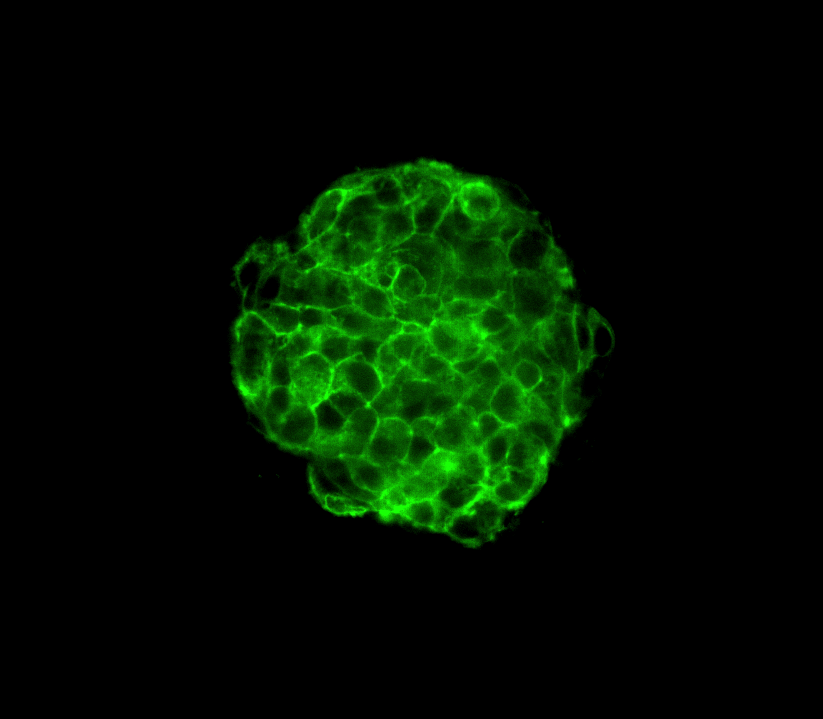

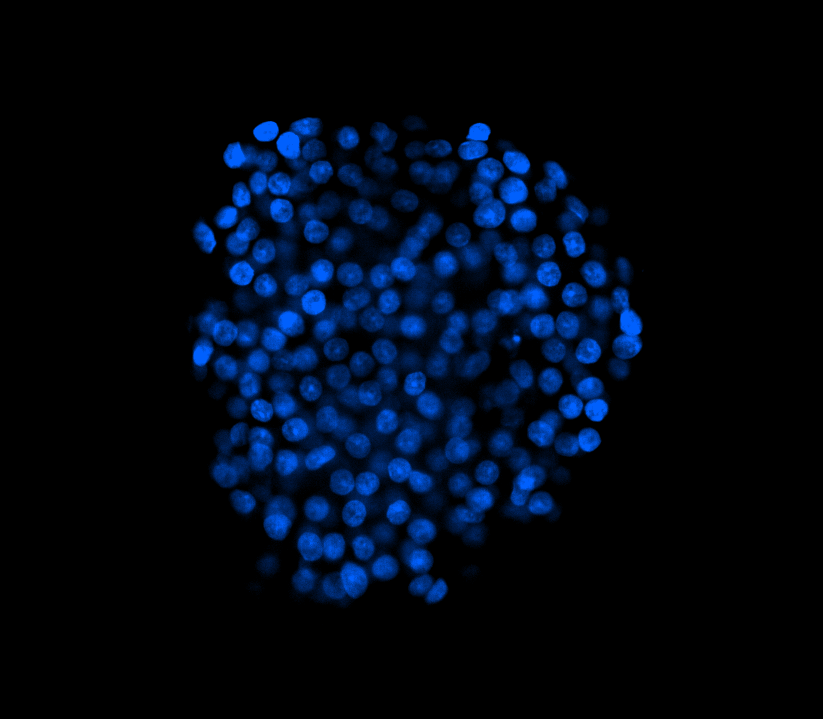

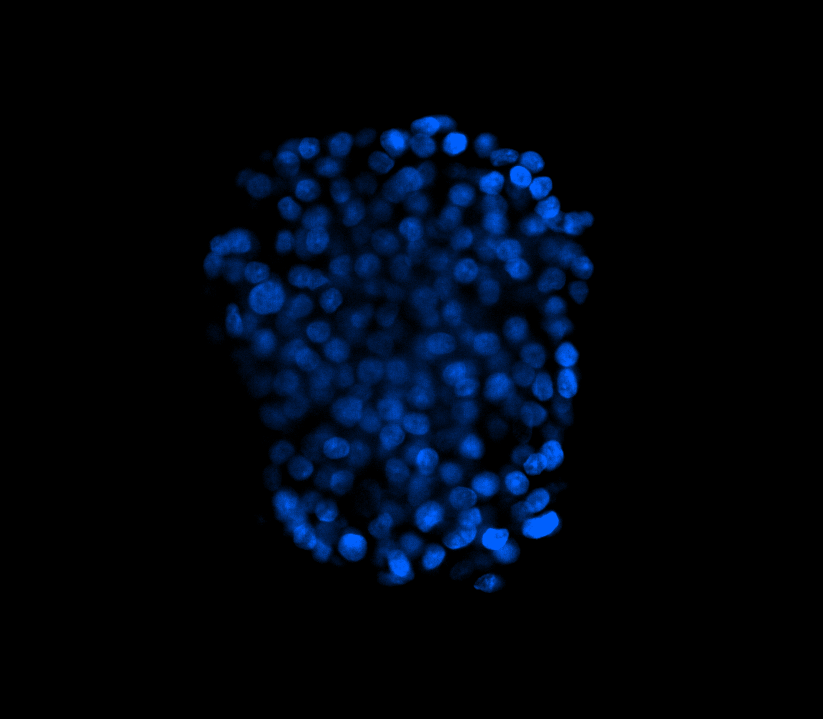

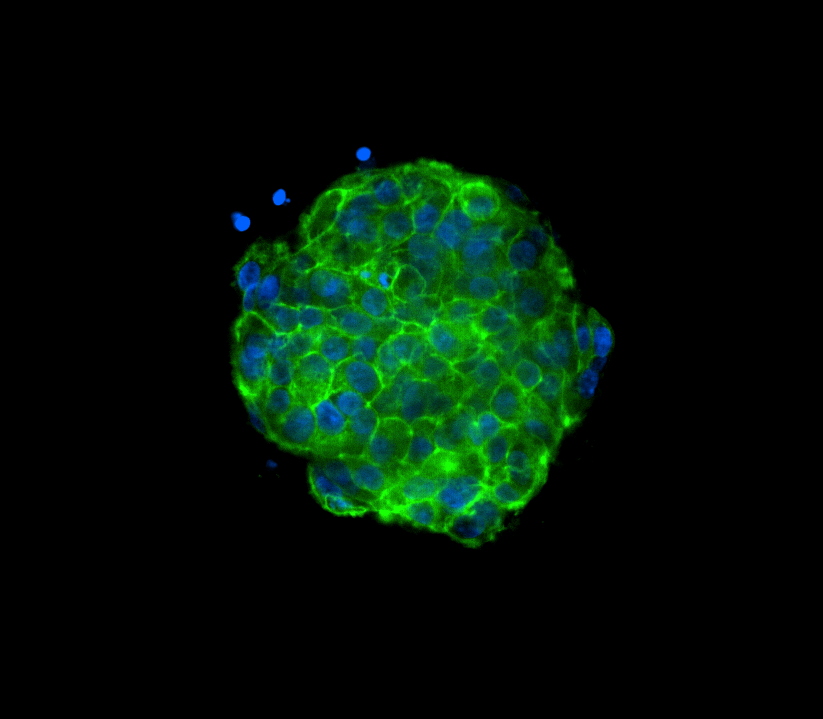

HLA-I

HLA-I

&

DAPI

shCTRL

shMEG3

Untreated

IFN-γ+IL-1β

+TNF-α +IFN-α 6d

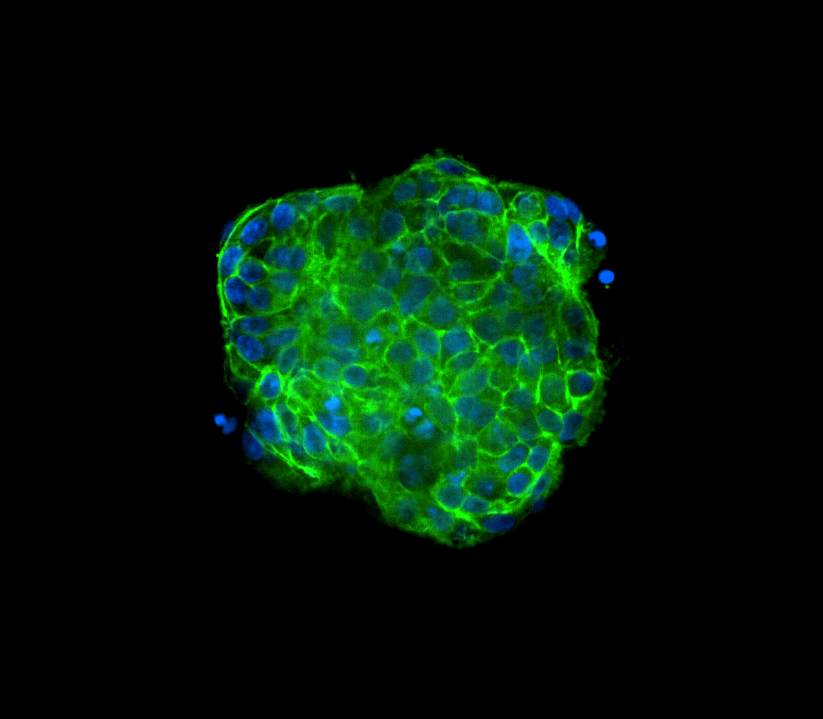

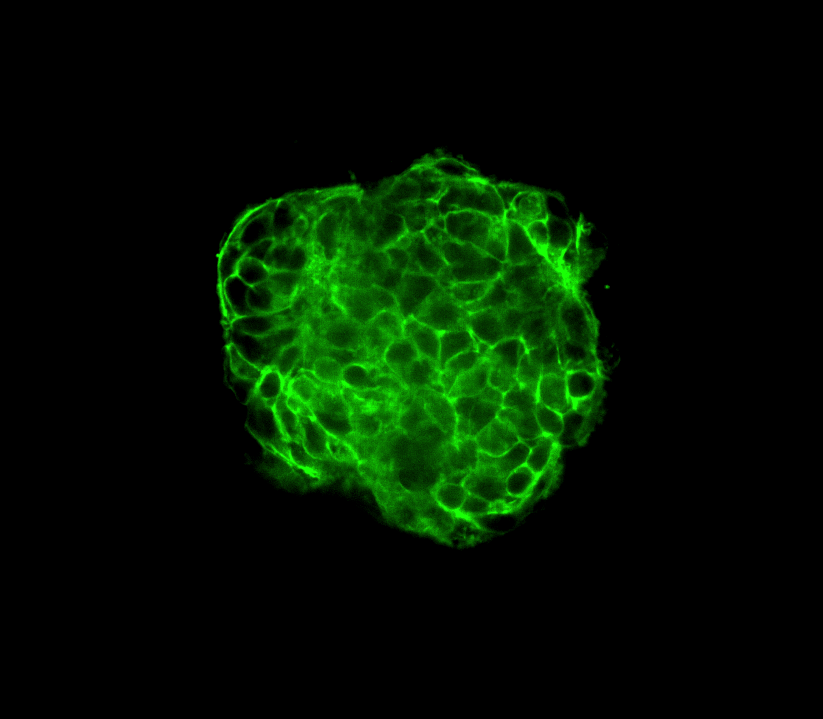

shCTRL

shMEG3

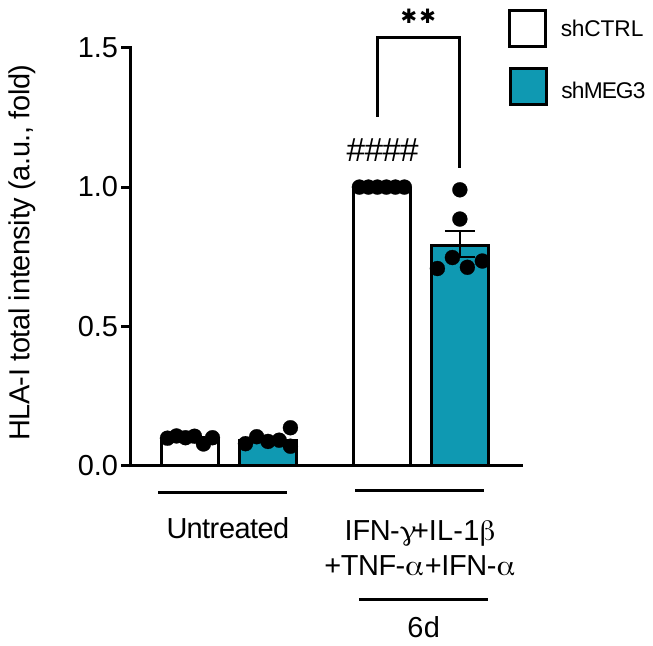

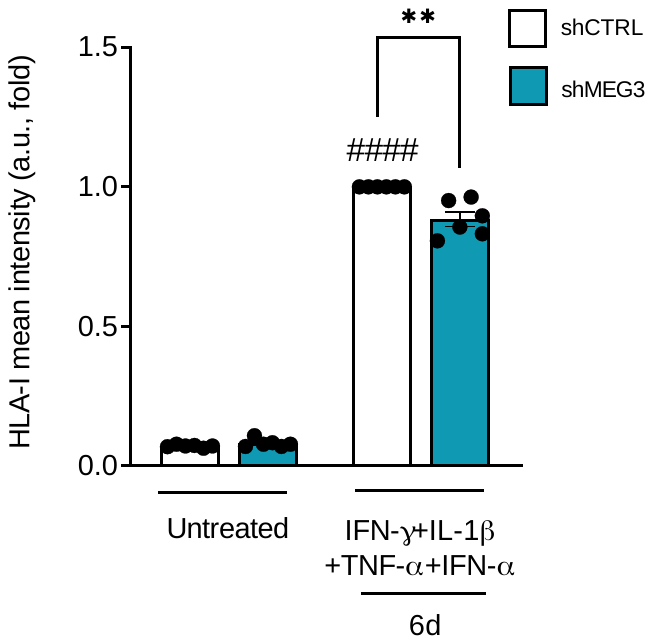

HLA-I

HLA-I

&

DAPI

A

B

C

**Figure S11. Silencing *MEG3* decreases cytokines-induced HLA-I expression in the cell surface in hIsMTs.** HIsMTs were transduced with short-hairpin RNA (shCTRL or shMEG3). After recovery, cells were left untreated or exposed to IFN-γ (25ng/mL) +IL-1β (5ng/mL) +TNF-α (25ng/mL) +IFN-α (10ng/mL) for 6 d. **(A)** Representative 3D immunofluorescence pictures of hIsMTs silenced for *MEG3* and exposed or not to cytokines stained for HLA-I (green). Nuclei were stained with DAPI (blue). **(B, C)** Total and mean intensities were determined using customized CellPathfinder pipelines. Results are means ± SEM. Each point represents a technical replicate. **p<0.01 vs shCTRL, ####p<0.0001 vs untreated one-way ANOVA, Bonferroni correction. HIsMTs: Human islets microtissues, SEM: Standard Error of the Mean, A.U.: arbitrary units.

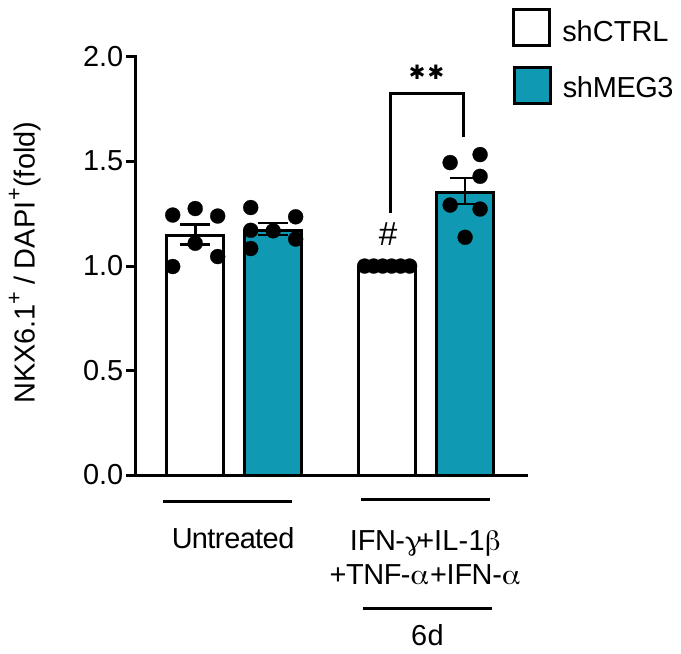

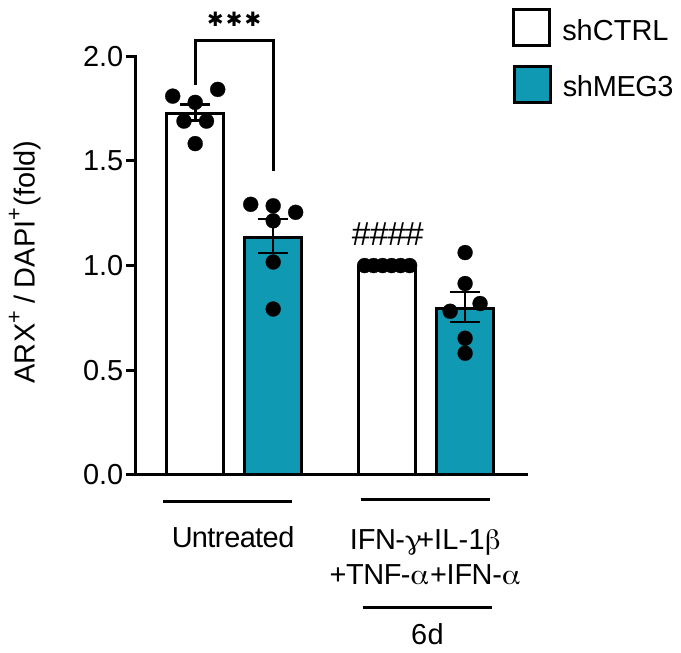

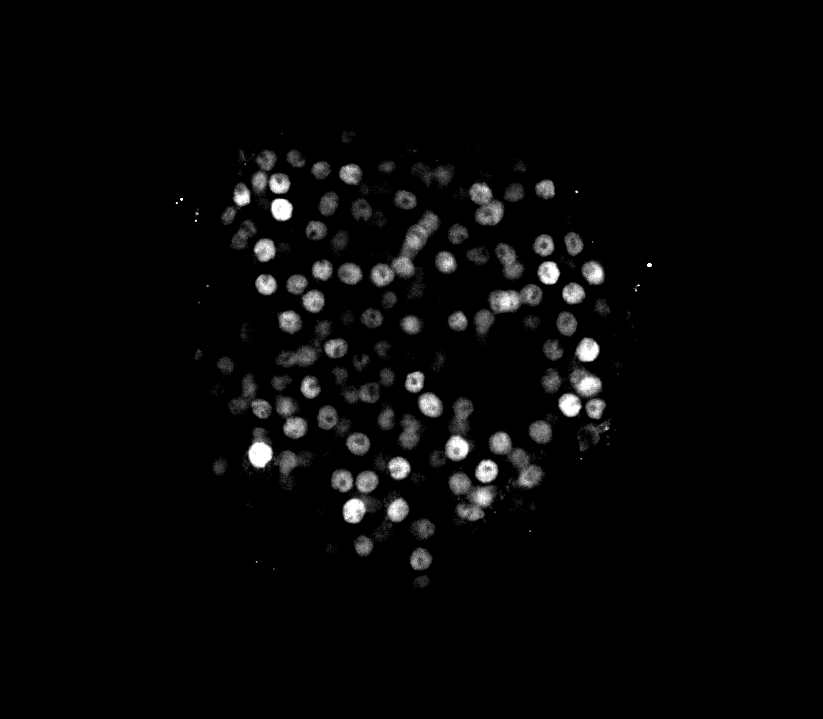

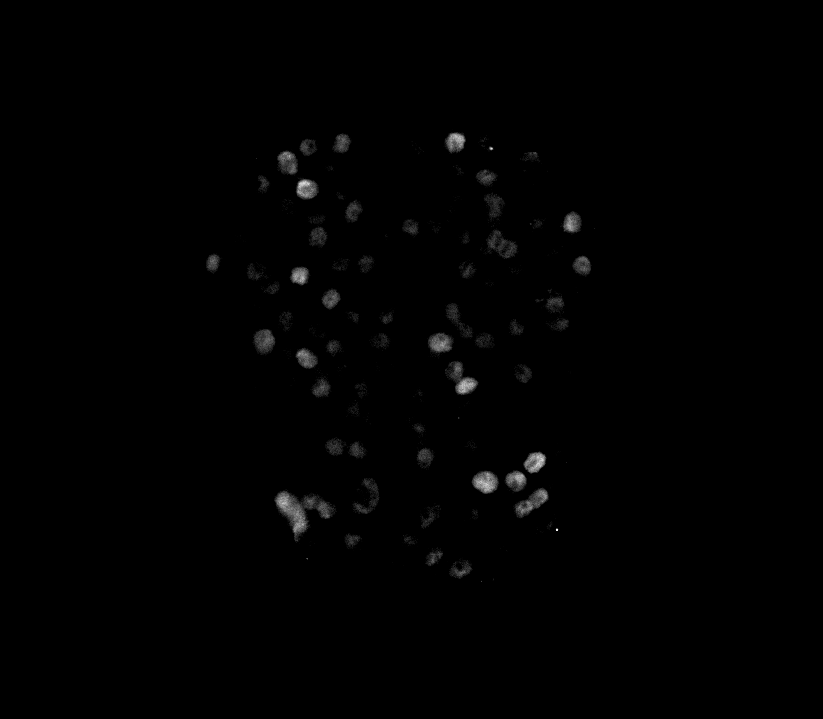

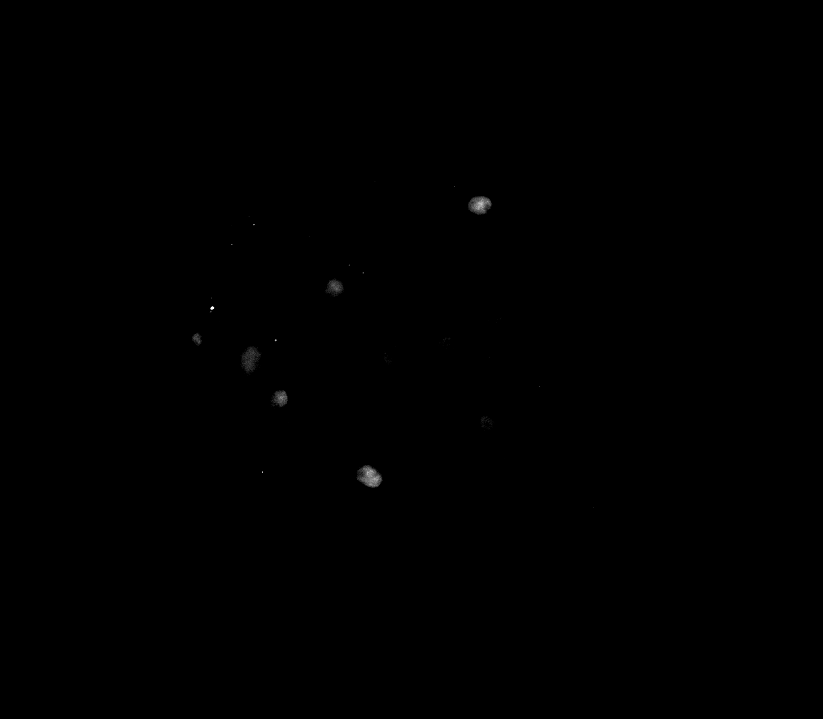

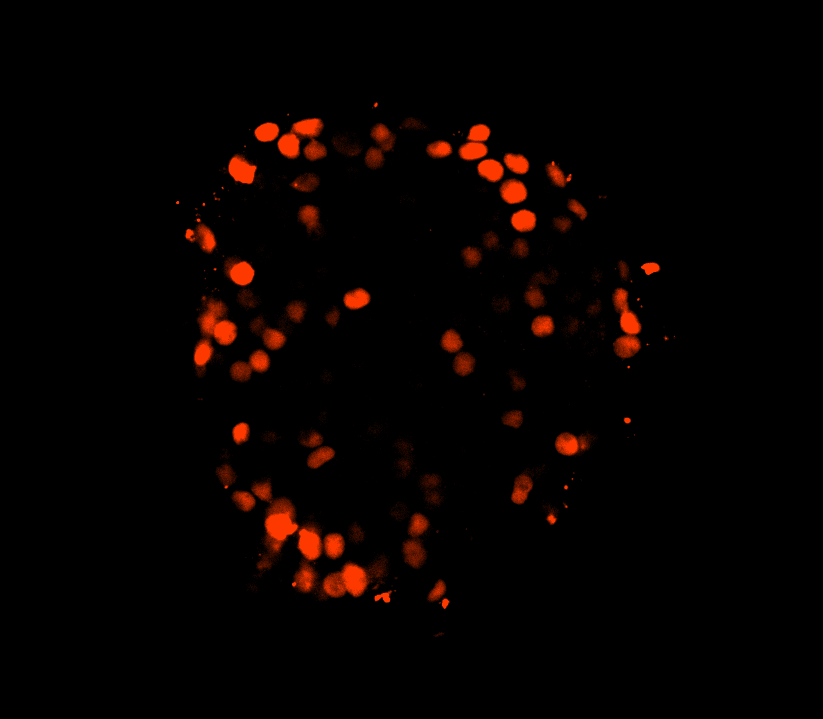

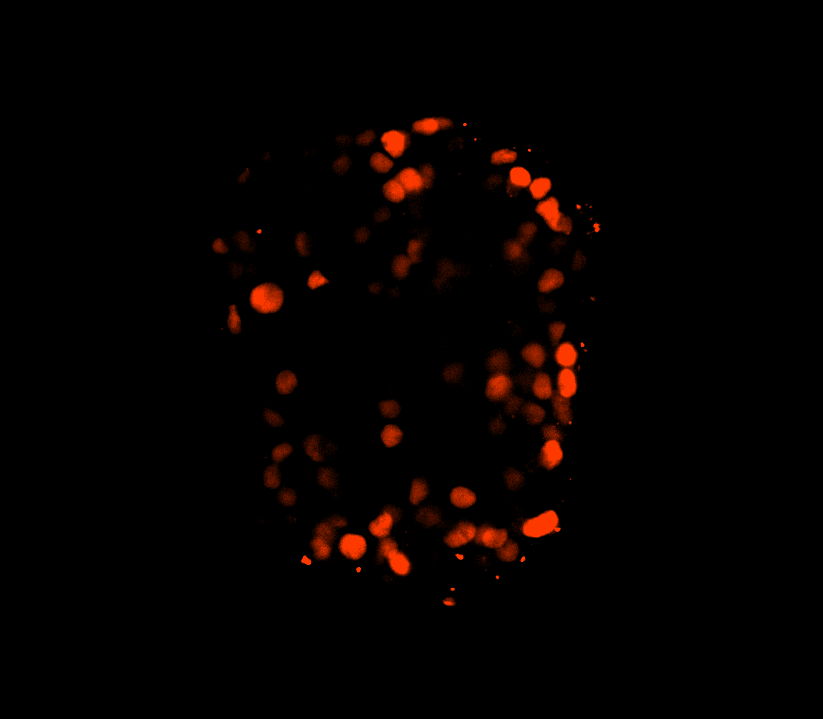

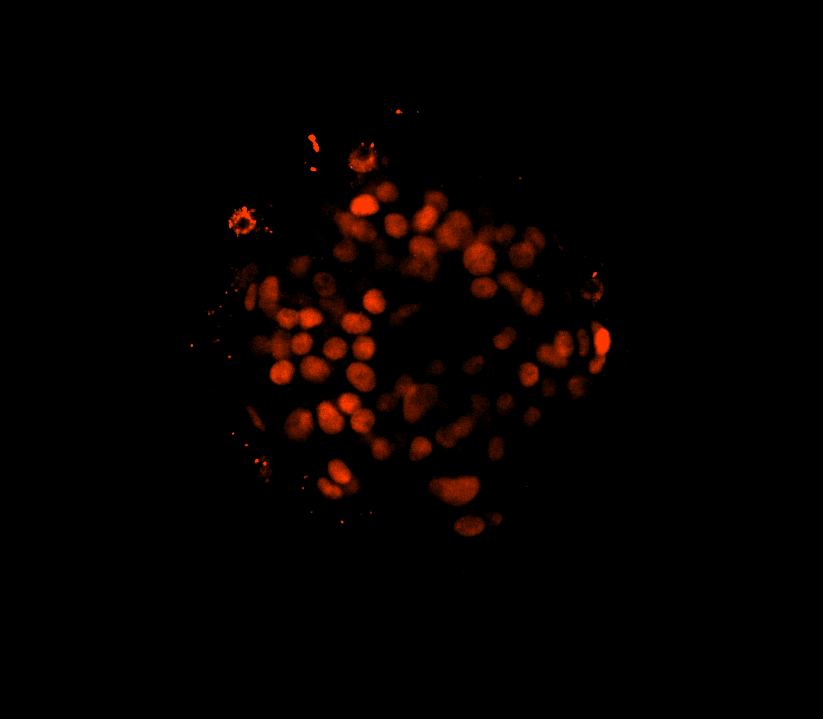

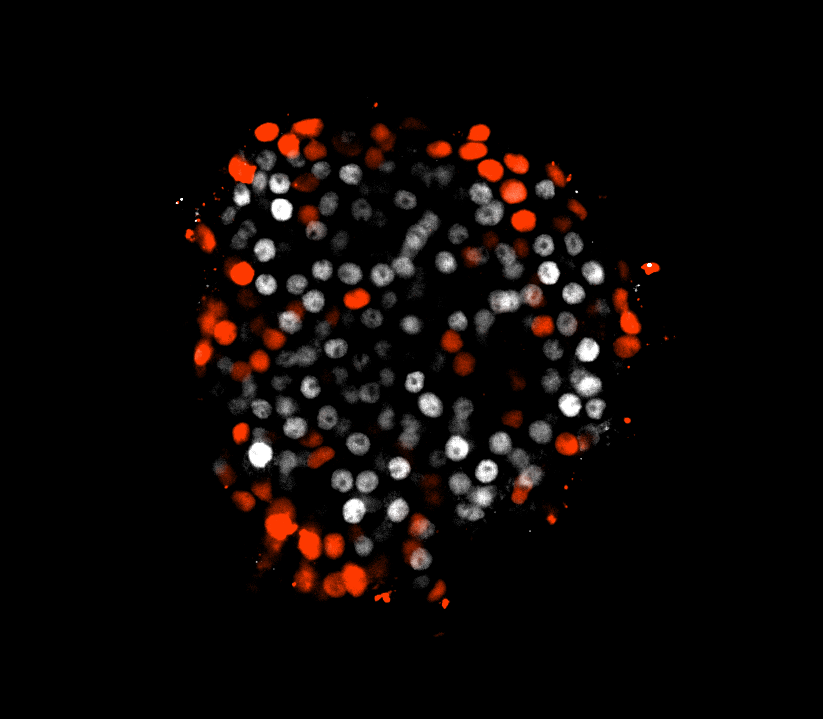

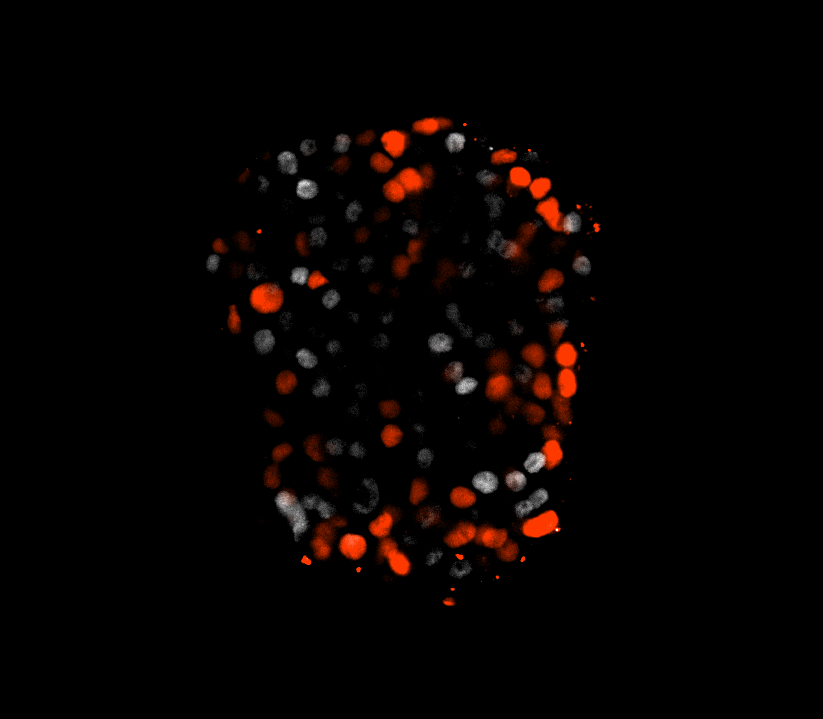

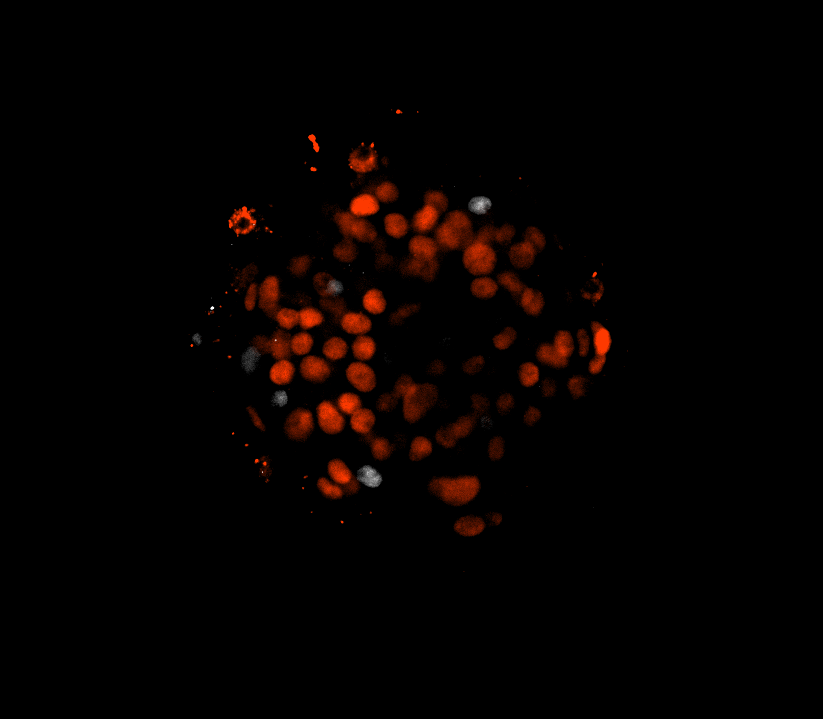

ARX

NKX6.1

ARX

&

NKX6.1

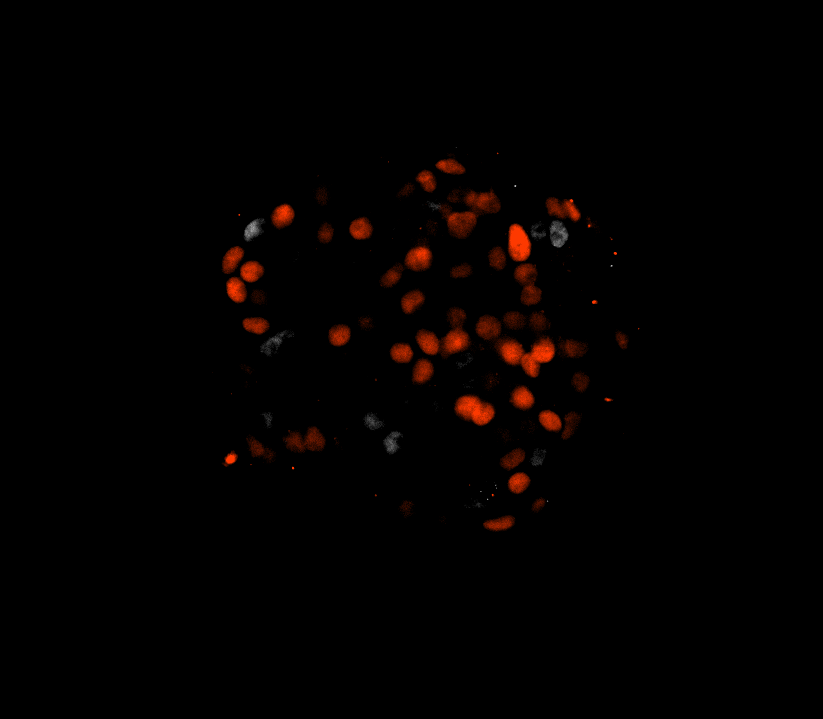

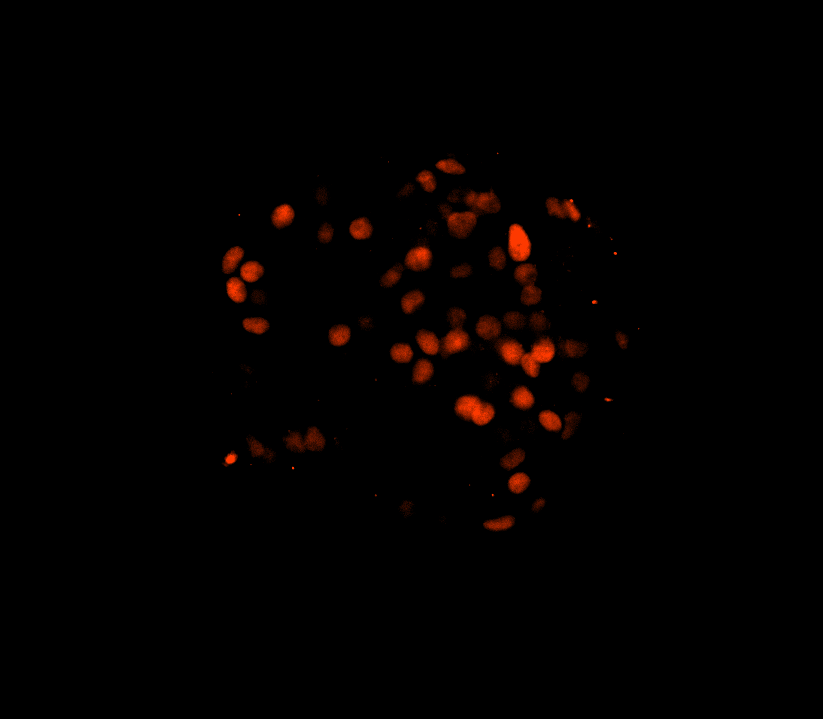

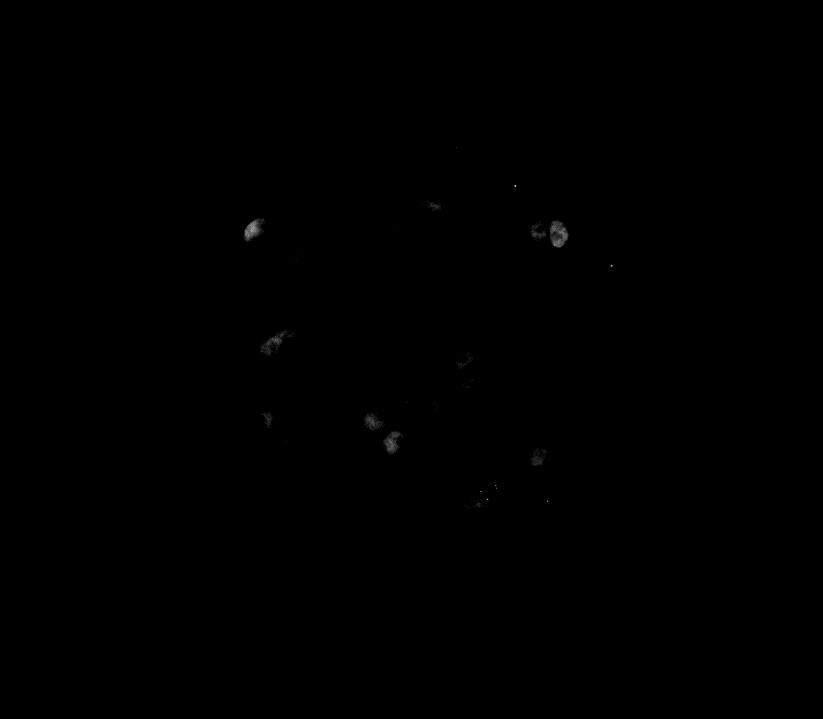

shCTRL

shMEG3

Untreated

IFN-γ+IL-1β

+TNF-α +IFN-α 6d

shCTRL

shMEG3

A

B

C

**Figure S12. Silencing *MEG3* modifies the proportion of alpha-/beta-cell fractions in hIsMTs.** HIsMTs were transduced with short-hairpin RNA (shCTRL or shMEG3). After recovery, cells were left untreated or exposed to IFN-γ (25ng/mL) +IL-1β (5ng/mL) +TNF-α (25ng/mL) +IFN-α (10ng/mL) for 6 d. **(A)** Representative 3D immunofluorescence pictures of hIsMTs silenced for *MEG3* and exposed to cytokines stained for NKX6.1 (red) and ARX (white). **(B, C)** Quantifications of **(B)** beta- and **(C)** alpha-cell fractions were performed using customized CellPathfinder pipelines. Results are means ± SEM. Each point represents a technical replicate. **p<0.01, ***p<0.001 vs shCTRL, #p<0.05, ####p<0.0001 vs untreated one-way ANOVA, Bonferroni correction. HIsMTs: Human islets microtissues, SEM: Standard Error of the Mean, A.U.: arbitrary units.

| **Dataset** | **ID** | **Description** | **Data type** | **Tissue** | **Samples** | **Age** | **Source** | **Year** |
| --- | --- | --- | --- | --- | --- | --- | --- | --- |
| 1 | HPAP-young (T1D control) | HPAP-young (T1D control) | scRNA-seq | Pancreatic islets | 15  (7F/8M) | 22.7  ±12.8 | HPAP [1] | 2025; 2022 |
| 2 | HPAP-AAB1+ | HPAP-AAB1+ | scRNA-seq | Pancreatic islets | 9  (3F/6M) | 43.1  ±10.5 | HPAP [1] | 2025; 2022 |
| 3 | HPAP-AAB2+ | HPAP-AAB2+ | scRNA-seq | Pancreatic islets | 2  (2M) | 48±15.6 | HPAP [1] | 2025; 2022 |
| 4 | HPAP-T1D | HPAP-T1D | scRNA-seq | Pancreatic islets | 11  (6F/5M) | 15.9±5.8 | HPAP [1] | 2025; 2022 |
| 5 | HPAP-older (T2D control) | HPAP-older (T2D control) | scRNA-seq | Pancreatic islets | 13  (7F/6M) | 42.5  ±10.7 | HPAP [1] | 2025; 2022 |
| 6 | HPAP-T2D | HPAP-T2D | scRNA-seq | Pancreatic islets | 10  (6F/4M) | 46.3±  8 | HPAP [1] | 2025; 2022 |
| 7 | Szymczak et al. | iPSC stage 7 | scRNA-seq | hiPSC-derived islet like cells | 5 | / | GSE203384 [2] | 2022 |
| 8 | Chandra et al. | iPSC stage 6 | scRNA-seq | hiPSC-derived islet like cells | 1 | / | GSE190726 [3] | 2022 |
| 9 | Sintov et al. (*in vivo*) | hESC *in vivo* | scRNA-seq | hESC-derived islet like cells | 5 | / | GSE200083 [4] | 2022 |
| 10 | Sintov et al. (*in vitro*) | hESC *in vitro* | scRNA-seq | hESC-derived islet like cells | 3 | / | GSE200084 [4] | 2022 |
| 11 | HPAP-basal | HPAP-basal | Bulk RNA-seq | FACS-sorted alpha-and beta cells | 12  (4F/8M) | 35.3 ±10.8 | HPAP [1] | 2024 |

**Table S1. Overview of the RNA-seq metadata for the analysed alpha and beta cells.** RNA-seq data of basal alpha- and beta cells were gathered from the Human Pancreas Analysis Program (HPAP, https://hpap.pmacs.upenn.edu/) and Gene Expression Omnibus (GEO) portal. Age is displayed as mean ± SD (when available for the donors). F: female, M: male, hiPSC: human induced pluripotent stem cells. hESC: human embryonic stem cells. T1D: type 1 diabetes, T2D: type 2 diabetes. AAB1+: donors positive for one islet autoantibody, AAB2+: donors positive for two or more islet autoantibodies.
