## Supplemental M&M for "Differential immune- and apoptosis-related gene signatures in pancreatic alpha and beta cells contribute to their fate in type 1 diabetes"

**scRNA-seq data processing additional steps**

Following the steps outlined in Material and Methods, "scRNA-seq data processing", we obtained high-quality single-cell data for downstream analysis. Gene expression normalization was performed using the SCTransform function in Seurat, which adjusts for variation in sequencing depth across cells. In addition, mitochondrial gene content was regressed out, as it reflects differences in cell quality or state. We identified the top 3,000 highly variable genes for principal component analysis (PCA). To integrate samples, we applied Harmony v1.2.0 using the top 50 principal components, correcting for sample origin and reagent kit batch effects as confounding variables. The resulting harmonized components were used to construct a Uniform Manifold Approximation and Projection (UMAP) embedding. Cell types were annotated using scSorter v0.0.2, based on known marker genes, and annotations were further verified by manual inspection of marker gene expression across clusters.

In this study, we focused on endocrine alpha and beta cells and therefore extracted these populations from all six datasets (**Figure S1**). To better visualize the final alpha and beta cell populations, we performed an Elbow plot analysis and selected 10 principal components (PCs), which captured ~80% of the total variance and the major sources of biological variation. These PCs were then used to generate UMAP embedding including only alpha and beta cells. For differential expression analysis between alpha and beta cells, we used the Wilcoxon rank-sum test, applied independently to each dataset. P-values were adjusted using Bonferroni correction based on the total number of detected genes. Genes with an adjusted P-value <0.05 were considered differentially expressed.

**Bulk RNA-seq analysis processing**

All samples underwent quality control using fastp v0.19.6 (1)*,* which performed adapter trimming, low-quality base filtering, and removal of reads shorter than 50 bp (i.e., less than half of the original 100 bp read length). To ensure high data quality and minimize potential confounding factors, we selected only samples with at least 30 million clean reads. We further restricted the analysis to donors who had both alpha cell and beta cell samples available from the same individual. Based on these criteria, 12 donors were included in the downstream analysis: HPAP-006, HPAP-014, HPAP-019, HPAP-022, HPAP-026, HPAP-035, HPAP-040, HPAP-052, HPAP-053, HPAP-054, HPAP-056, and HPAP-080 (**Table S1**). Clean reads were aligned to the human transcriptome reference (GENCODE v36, GRCh38.p13) using Salmon v1.4.0 (2), with the parameters “--seqBias, --gcBias, and –validateMappings” enabled to correct for sequencing and GC content bias. The Salmon index was generated using default parameters and k-mer settings. Transcript-level quantifications after alignment were imported using the tximport package (3) and summarized to gene-level counts via the summarizeToGene function. Differential gene expression analysis was performed with DESeq2 v1.38.3 (4). Dispersion estimates and log2 fold changes were computed using the Wald test, and Benjamini-Hochberg correction was applied to control the false discovery rate. Genes with an adjusted P-value < 0.05 were considered differentially expressed.

**Rank-rank Hypergeometric Overlap (RRHO) analysis**

Using the enhanced RRHO pipeline RedRibbon (5), genes detected in both datasets were independently ranked by log2 fold change - derived from differential gene expression analysis of scRNA-seq data - along the x- and y- axis, respectively. For each coordinate in the rank space, RedRibbon applies a hypergeometric test to evaluate the statistical significance of overlapping genes in the corresponding rank windows. RedRibbon employs an evolutionary algorithm to explore the rank space and uses permutation-based adjustment to calculate significance values for each gene rank combination, generating a level map that visualizes -log10 P-values across the gene rank space. The resulting heatmap highlights four regions of transcriptional concordance or discordance: up–up, down–down, up–down, and down–up. As a reference, we generated a simulated input data matrix in which the log2 fold change values were identical, resulting in a perfect correlation (or concordance) map (**Figure S3B**).

**Interferon-stimulated gene (ISG) score calculation**

Based on our previous transcriptomic analysis (6), we defined a beta cell-specific interferon-stimulated gene (ISG) signature by identifying mRNAs that were robustly (>3-fold) upregulated in insulin-producing EndoC-βH1 cells in response to IFN-α or IFN-γ stimulation but remained unaffected by IL-1β or TNF-α exposure. The list of genes in the ISG signature consists of 53 genes, including key regulators such as interferon regulated factors (*IRF1, IRF2, IRF7, IRF8* and *IRF9*) and STAT transcription factors (*STAT1*, *STAT2* and *STAT3*), various HLA class I and II genes (*HLA-A, HLA-B, HLA-C, HLA-E, HLA-F, HLA-DRA, HLA-DMB* and *HLA-DQB1*), chemokines (*CXCL8, CXCL9, CXCL10, CXCL11, CXCL14* and *CXCL17*), interleukins and related receptors (*IL12A*, *IL15RA, IL18BP* and *IL6*), and other immune effectors (*NLRC5, PSMB10, PSMB8, PSMB9, TNFRSF1B, CASP1, RIG-I, DDX60, IDO1, IFI16, MDA5, MX1, OAS1, OAS3, PNPT1, TREX1, ZBP1, ZNFX1, SOCS1, SOCS3, USP18, C1QB, C4A, CCL14, CIITA* and *CSF1*). In the current study, ISG scores were calculated as the average transcript per million (TPM) expression of these 53 genes for each alpha- or beta-cell sample after fluorescence-activated cell sorting (FACS).

The pro-inflammatory and anti-inflammatory gene signatures were selected based on a previously published review from our lab (7) and consist of 15 and 16 genes, respectively. The pro-inflammatory gene signature includes: *MAPK8*, *MAPK9*, *TYK2*, *STAT1*, *STAT2*, *NFKB1*, *IRF1*, *IRF9*, *CXCL10*, *HLA-A*, *HLA-B*, *HLA-C*, *B2M*, *IRF7*, and *DDIT3*. The anti-inflammatory gene signatures include: *SOCS1*, *SOCS2*, *SOCS3*, *PTPN2*, *BACH2*, *NFKBIA*, *NFKBIB*, *USP18*, *ISG15*, *HLA-E*, *BCL2L1*, *IFIH1*, *IL10*, *CAT*, *SOD2*, and *HSPA5*. In the current study, pro- and anti-inflammation scores were calculated as the average expression of the respective gene sets at the single-cell level for each group of alpha and beta cells.

**Quantitative real-time PCR, protein extraction and Western blott analysis**

The lysates from hIsMTs were snap-frozen in liquid nitrogen and total RNA was extracted using the RNeasy Micro Kit (Qiagen) and reverse-transcribed using the Reverse Transcriptase Core Kit (Eurogentec). The quantitative RT-PCR amplification reactions were performed with IQ SYBR Green Supermix (Bio-Rad Laboratories) and ran in the CFX Connect Real-Time PCR Detection System (Bio-Rad Laboratories). The product quantification was performed using the standard curve method (8). For each gene, the melting curve was analysed to confirm amplification of a single PCR product. Gene expression values were normalized by the geometric mean of the housekeeping genes *ACTIN* and *VAPA* (9) and presented as fold change as detailed in the figure legends. The primers used are listed below: *ACTIN*; FW: CTGTACGCCAACACAGTGCT, RV: GCTCAGGAGGAGCAATGATC, *BiP;* quantiTect primer #QT00096404, *CHOP*; quantiTect primer #QT00082278, *CXCL10*; FW: GTGGCATTCAAGGAGTACCTC, RV: GCCTTCGATTCTGGATTCAG, *INSULIN*; FW: AGCCCTCCAGGACAGGC, RV: TTTGCTGGTTCAAGGGCTTT, *MEG3*; FW: CCCTCTTGCTTGTCTTACTTGTC, RV: CCTTTCAAGAAGCTTGGCTGATG, *PDX1*; FW: AAAGCTCACGCGTGGAAA, RV: GCCGTGAGATGTACTTGTTGA, *VAPA*; FW: TACCGAAACAAGGAAACTAATGGAA, RV: GCCTTAAACCTTCATCTCTCAGGT.

**3D staining and imaging**

After fixation, as described in Material and Methods, hIsMTs were permeabilized with permeabilization buffer (Triton® X-100, 0.5% in PBS w/o Mg_2_^+^Ca_2_^+^) and washed twice with PBS. hIsMTs were blocked with 5% donkey serum in PBS to prevent nonspecific antibody binding. hIsMTs were then incubated overnight with primary antibody: rabbit anti-NKX6.1 (Abcam, ab221549); sheep anti-ARX (R&D Systems, AF7068-SP); mouse anti-HLA-ABC (HLA-I, Biolegend, 311427) in antibody dilution buffer (5% donkey serum, 0.2% Triton® X-100 in PBS w/o Mg_2_^+^Ca_2_^+^). hIsMTs were washed with wash buffer (0.2% Triton® X-100 in PBS w/o Mg_2_^+^Ca_2_^+^) and incubated with secondary antibodies (donkey anti-rabbit AF568 (Thermofisher, A10042); donkey anti-sheep AF647 (Jackson ImmunoResearch, 713-605-147) in antibody dilution buffer, along with DAPI (D9542). hIsMTs were washed 3 times with wash buffer, transferred into Akura 384-well ImagePro plates (InSphero, CS-PC14), and cleared with ScaleS4 solution for 3D imaging. Images were acquired using Yokogawa CQ1 Benchtop High-Content Analysis System, taking fluorescent images in 3 µm z-steps. Images were quantified using customized CellPathfinder pipelines.
